# A physiology-based mathematical model of the renin-angiotensin system, bone remodeling, and calcium homeostasis: Effects of estrogen and renin-angiotensin system inhibitors

**DOI:** 10.1101/2025.05.01.651663

**Authors:** Melissa M. Stadt, Anita T. Layton

**Affiliations:** Department of Applied Mathematics, University of Waterloo, Waterloo, ON, Canada; Cheriton School of Computer Science, Department of Biology, School of Pharmacology, University of Waterloo, Waterloo, ON, Canada

## Abstract

During menopause, estrogen levels decline significantly, leading to substantial physiological changes due to estrogen’s regulatory role in various systems. In particular, estrogen helps prevent excessive bone resorption by its impact on bone remodeling. When estrogen levels decrease, bone resorption increases, often resulting in weakened bones and osteoporosis in post-menopausal women. Experimental studies have also shown that estrogen regulates the renin-angiotensin system (RAS), a hormone system involved in many physiological processes, including blood pressure regulation. Additionally, the RAS has an interconnected relationship with calcium regulatory and bone remodeling systems. Given these dynamic interplays, how would perturbations in one system affect the others? To answer that question, we developed a physiology-based mathematical model that simulates the interactions of estrogen, key RAS components, calcium regulation, and bone remodeling. Through sensitivity analysis and model simulations, we investigated how declining estrogen levels affect the RAS and bone mineral density. Furthermore, we quantified how RAS inhibitors, specifically angiotensin-converting enzyme inhibitors and angiotensin receptor blockers, may increase bone density during post-menopausal estrogen decline.

## 1 Introduction

Bone remodeling is a complex physiological process that maintains the balance between bone formation and resorption. This process is mediated by bone cells: osteoblasts, which are responsible for bone formation, and osteoclasts, which are responsible for bone resorption. The activity and proliferation of osteoblasts and osteoclasts is regulated by a system of various receptors, their corresponding ligands, and hormonal signals [1, 2]. Estrogen, in particular, impacts bone remodeling by inhibiting osteoclast activity, thereby protecting bones from excessive resorption. When estrogen levels in women decline post-menopause, osteoclast activity increases, resulting in increased resorption, and ultimately weakening bones. This often leads to post-menopausal osteoporosis. Post-menopausal osteoporosis is the most common form of osteoporosis, which is characterized by increased bone fragility, making fractures more likely from minor trauma or in severe cases, even spontaneous fracture. It is estimated that about 1 in 3 women over 50 years of age will suffer a fracture related to post-menopausal osteoporosis [3, 4].

The main component of bone structure is hydroxyapatite, a compound made up of phosphate and calcium. Given their role in developing the key structure of bones, maintaining balanced levels of calcium and phosphate is critical for bone health. The regulation of calcium and phosphate levels is driven by calciotropic hormones: calcitriol and parathyroid hormone (PTH) [5]. Researchers have reported a bidirectional relationship between the calciotropic hormones and the renin-angiotensin system (RAS) [6], a hormone system that plays a key role in many physiological processes, including blood pressure regulation, in addition to a direct impact of the RAS on bone remodeling [7]. The RAS is a hormone cascade that produces the bioactive peptide angiotensin II (Ang II). Ang II binds to angiotensin type 1 receptors (AT1R) and angiotensin type 2 receptors (AT2R) to trigger various physiological effects. The RAS is also modulated by estrogen levels, and therefore is impacted by the rapidly declining estrogen levels in post-menopausal women [8]. This dysregulation contributes to the higher susceptibility of post-menopausal women to the development of hypertension, cardiovascular disease, and osteoporosis [8].

Inhibition of RAS components, particularly angiotensin-converting enzyme (ACE) and AT1R binding, has shown to be effective targets for antihypertensive treatment [9]. These drugs are classified as RAS inhibitors. Experimental data has shown that the use of RAS inhibitors improves bone quality independent of their effect on blood pressure [10, 11]. Given that RAS inhibitors are relatively well tolerated, safe, and are reasonably priced drugs, there has been interest in using RAS inhibitors to prevent osteoporosis in post-menopausal women [10–12]. Additionally, since hypertension is highly prevalent in post-menopausal women, understanding the impacts of antihypertensive treatments, such as RAS inhibitors, on bone remodeling is important for achieving blood pressure control and bone health in post-menopausal women [10–12].

Many mathematical models have been developed to study bone remodeling as recently reviewed in Ref. [2]. Peterson and Riggs [13] pioneered a mathematical model of calcium homeostasis coupled to a detailed bone biology model. Their model consists of bone, renal, and blood compartments to predict levels of PTH, calcitriol, calcium, phosphate, and bone remodeling activity. This model has been applied to investigate applications related to bone remodeling and calcium homeostasis, including the impact of estrogen [14], mineral bone disorder in patients with chronic kidney disease [15], and the impact of fibroblast growth factor 23 and vascular calcification [16]. Mathematical models have also been developed for the RAS to understand its role in blood pressure regulation. Specifically, Lo et al. [17] developed a model of the RAS in humans to investigate the differences between normo-tensive and hypertensive patients. This model was later included in a full body blood pressure model by Hallow et al. [18]. Leete et al. [19] further developed this model to investigate the effects of sex-based differences in the RAS on blood pressure regulation. However, no known models have been developed that include the direct impact of estrogen on the RAS or have coupled the RAS impacts with calcium homeostasis and bone remodeling in humans.

In this study, we (i) investigate the impact of age-related estrogen decline by including the impact of estrogen on the RAS and (ii) couple the calcium homeostasis and bone remodeling model (Refs. [13, 14]) with the RAS model (Refs. [17–19]) to predict the impact of RAS inhibitors on this system.

## 2 Methods

In this study, we develop a coupled ordinary differential equation model that represents (i) the impact of estrogen on the RAS (Section 2.1.1) and (ii) couple this estrogen-RAS model with a bone biology and calcium homeostasis model (Section 2.3). We then use this model to simulate the effects of post-menopausal estrogen decline on the RAS, bone remodeling, and calcium homeostasis. Additionally, we simulate the impact of RAS inhibitors. To quantify the impact of parameter variation on model output, we conduct a global sensitivity analysis. A model schematic is shown in Fig. 2.1. A list of the model equations is given in Appendix D.

**Figure 2.1:**
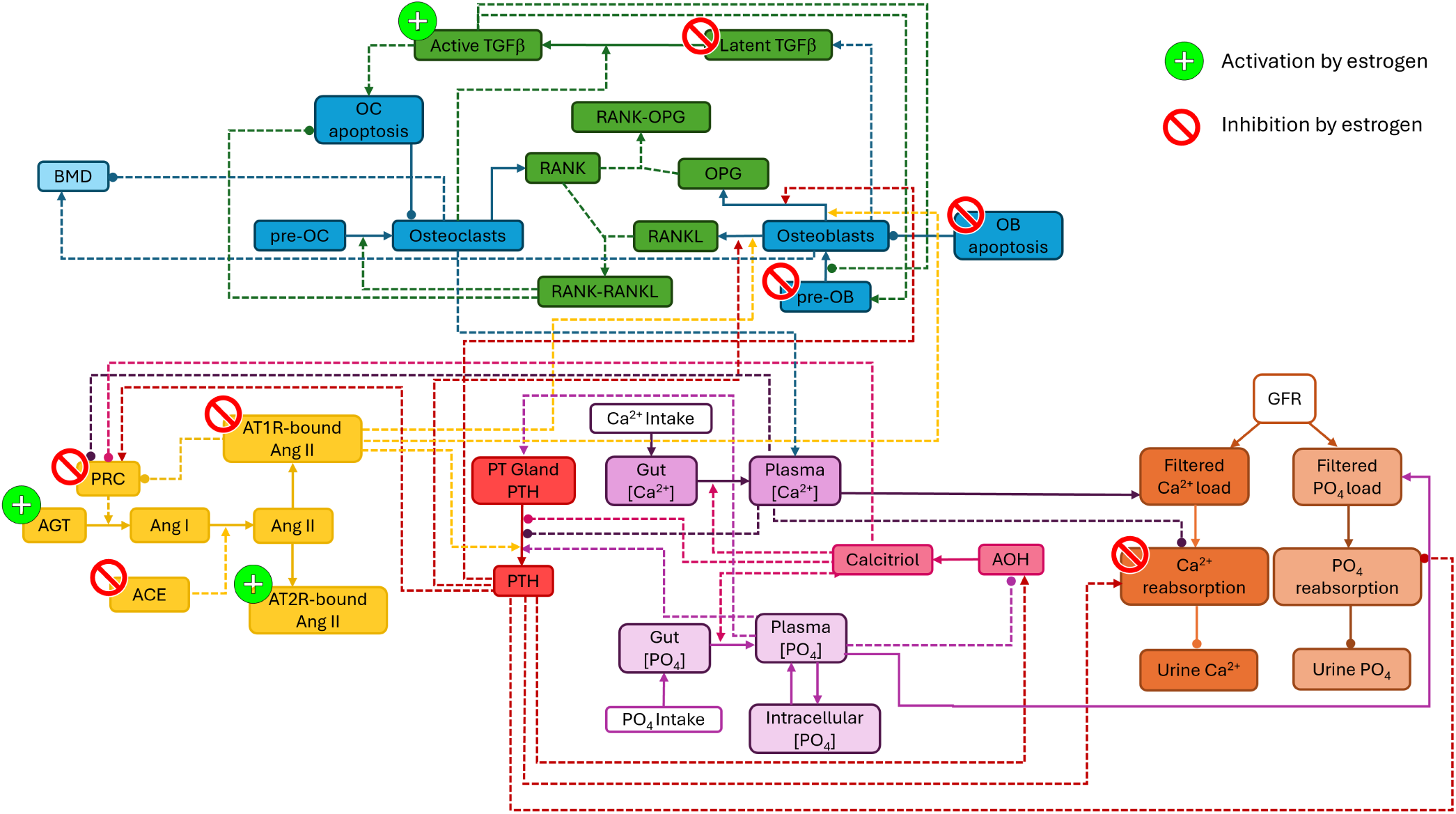
Schematic model of calcium regulation, bone remodeling, the renin-angiotensin system (RAS), and the impact of estrogen on these systems. Purple nodes represent phosphate and calcium levels, orange nodes represent phosphate and calcium renal handling. Yellow nodes represent the RAS, red nodes represent parathyroid hormone, pink nodes represent calcitriol, blue nodes represent bone cells and bone mineral density, green nodes represent signaling mechanisms in the bone. Nodes with a white background represent fixed values in the model. The impacts of estrogen are indicated by the red and green symbols for inhibition and activation, respectively. The solid arrows represent flux, secretion, synthesis, or expression, while the dashed lines represent impacts on other biological components. The pointed arrows represent activation and the rounded arrows represent inhibition. ACE: angiotensin-converting enzyme, AGT: angiotensinogen, Ang: angiotensin, AOH: 1α-hydroxylase, AT1R: angiotensin type 1 receptor, AT2R: angiotensin type 2 receptor, BMD: bone mineral density, Ca^2+^: calcium, GFR: glomerular filtration rate, OB: osteoblast, OC: osteoclast, OPG: osteoprotegerin, PO_4_: phosphate, PRC: plasma renin concentration, PT: parathyroid, PTH: parathyroid hormone, RANK: Receptor Activator of Nuclear Factor Kappa-B, RANKL: RANK ligand, TGFβ: transforming growth factor β

### 2.1 Renin-angiotensin system model

In this study, we use the RAS model with the female-specific parameter set from Ref. [19]. The model variables consist of the systemic concentrations of angiotensinogen (AGT), Ang I, Ang IV, Ang (1-7), AT1R-bound Ang II, and AT2R-bound Ang II. AT1R-bound Ang II inhibits renin secretion through a feedback loop. For a detailed description of the underlying RAS model used in this study, see Refs. [18, 19].

#### 2.1.1 Impact of estrogen decline in aging women on the RAS

Estrogen levels impact the RAS by activating and inhibiting various key processes. Specifically, estrogen activates AGT production and AT2R binding, while inhibiting renin secretion, ACE activity, and AT1R binding [8]. These processes are illustrated in Fig. 2.1.

After the onset of menopause, estrogen levels decrease dramatically. To model this decline in estrogen levels, we use the estrogen model for post-menopausal women from Ref. [20]. The normalized level of estrogen, denoted by *E*, is modeled through explicit time dependence so that

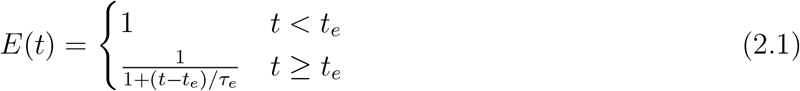

where *t* is the time in years (i.e., age), *t*_*e*_ = 50 years is the age of onset of estrogen decline, and *τ*_*e*_ = 2.6 years is a parameter fit to normalized estrogen data from Ref. [21] that determines the slope of the decline in estrogen (see Ref. [20] for details). Note that before *t*_*e*_, *E* = 1 so that estrogen is assumed to remain at baseline level pre-menopause. Indeed, estrogen levels vary during the menstrual cycle [22]. In this study, we do not consider variations in estrogen over the menstrual cycle in pre-menopausal women and consider baseline estrogen level pre-menopause as the average level through the full cycle.

To include the impact of estrogen on the RAS, we use functions of the form:

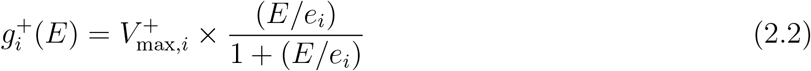

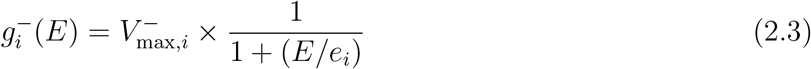

to model activation (*g*^+^) or inhibition (*g*^*−*^) with

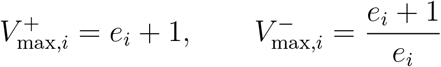

where *i* denotes the target mechanism, *e*_*i*_ is a parameter that is determined for each relevant target mechanism, and *E* is given by Eq. 2.1. Note that when 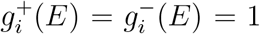 for any value of *e*_*i*_, so that at baseline estrogen levels, there are no changes.

The enzyme renin is secreted by the juxtaglomerular cells of the kidneys to convert AGT to Ang I. Estrogen inhibits renin levels [8, 23, 24]. We let 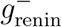 be the impact of estrogen on renin secretion so that renin secretion is modeled by

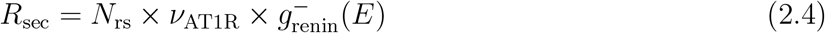

where *N*_rs_ is baseline renin secretion and *ν*_AT1R_ is the feedback of [AT1R-bound Ang II] on renin secretion from Ref. [18]. Plasma renin concentration (PRC) is modeled by

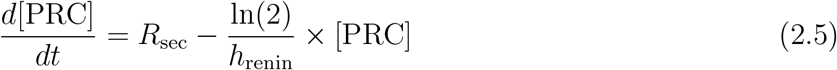

where *h*_renin_ denotes the half life of renin as in Ref. [18].

AGT is synthesized in the liver and is stimulated by estrogen [8, 23, 25]. To model the impact of estrogen on AGT, we multiply the production of AGT, denoted by *k*_AGT_, by 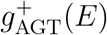 so that [AGT] is modeled by

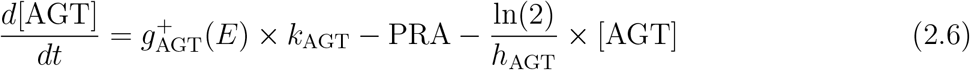

where

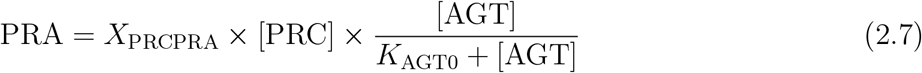

is plasma renin activity (PRA) with parameter *X*_PRCPRA_ and *h*_AGT_ is the half-life for AGT as in Ref. [18]. The equation for PRA is a slight modification from the equation used in Refs. [18, 19] to include the impact of AGT on PRA as well. The *K*_AGT0_ parameter was fit to predict the same model output in the original blood pressure model from Ref. [19]. In post-menopausal women, AGT levels decline from pre-menopause levels [23].

Angiotensin-converting enzyme (ACE) converts Ang I into Ang II. Researchers have reported that both ACE expression and activity are inhibited by estrogen in rodent models [26–29]. We let 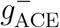 represent the inhibition of estrogen levels on ACE activity so that the concentration of Ang I and Ang II are modeled by

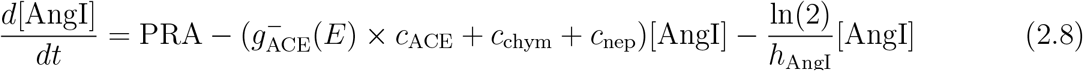

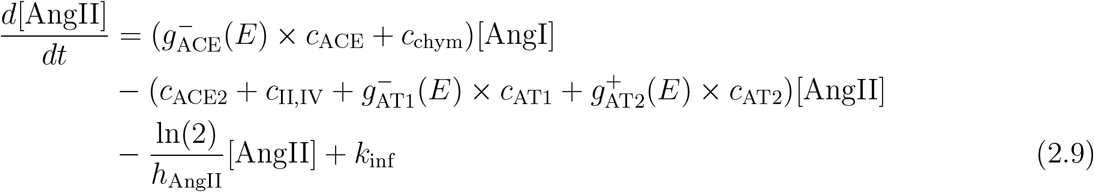

where *c*_*i*_ denotes the corresponding enzyme activities and *h*_AngI_ and *h*_AngII_ are the half lives of Ang I and Ang II, respectively, as in Ref. [18]. *k*_inf_ is an input used for Ang II infusion (see Section 2.3.4) and is set to 0 in simulations with no Ang II infusion.

Ang II is reported to remain around the same in aging humans [30]. Ovariectomized rats showed a small decrease in plasma Ang II levels, which was reversed by estrogen replacement at preovariectomy levels [31,32]. We note that Ang II levels are often reported in post-menopausal women on estrogen replacement therapy, but researchers have found significant differences in the impact of estrogen replacement therapy on the RAS when taking via oral or transdermal therapy [33]. This is likely due to the high exposure to estrogen in the liver when ingesting oral estrogen.

Ang II binds to AT1R and AT2R, triggering different physiological responses. Estrogen inhibits Ang II binding to AT1R, but activates Ang II binding to AT2R [8, 29, 34]. Angiotensin type 1 receptor density is increased over 2-fold in ovariectomized rats [35]. We based our parameter value for *e*_AT1_ on these results. AT1R-bound Ang II ([AT1R-AngII]) and AT2R-bound Ang II ([AT2R- AngII]) concentrations are modeled by

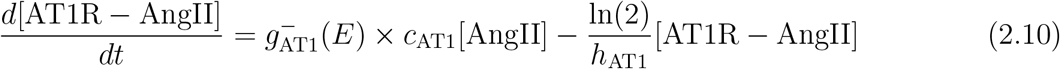

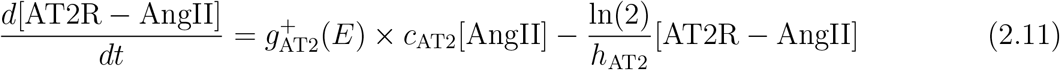

with *h*_AT1_ and *h*_AT2_ denoting the half-lives of AT1R-bound Ang II and AT2R-bound Ang II from Ref. [18]. *c*_AT1_ and *c*_AT2_ denote the binding rates of Ang II to AT1R and AT2R, respectively, as in Ref. [18]. 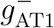 and 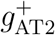 represent the impact of estrogen on the respective binding rate.

We chose parameter values for *e*_*i*_ based on reported changes in the respective RAS levels in the literature as described above. The values for *e*_*i*_ for each RAS component impacted by estrogen levels are listed in Table 2.1. Model simulations of the estrogen-RAS model in isolation are shown in Appendix A.

**Table 2.1:** Values of the parameters e_i_ that determine the impact of estrogen on renin-angiotensin system components. The e_i_ parameters are unitless.

| Name | Value | Equation(s) |
| --- | --- | --- |
| $e_{AGT}$ | 0.15 | $g_{AGT}^+$, Eq. 2.6 |
| $e_{renin}$ | 0.15 | $g_{renin}^-$, Eq. 2.4 |
| $e_{ACE}$ | 5.0 | $g_{ACE}^-$, Eqs. 2.8, 2.9 |
| $e_{AT1}$ | 0.5 | $g_{AT1}^-$, Eqs. 2.9, 2.10 |
| $e_{AT2}$ | 0.05 | $g_{AT2}^+$, Eqs. 2.9, 2.11 |

#### 2.1.2 RAS inhibitor simulations

To simulate the RAS inhibitors, we use the same approach as in Ref. [18]. Specifically, to simulate treatment with an angiotensin receptor blocker (ARB), we let

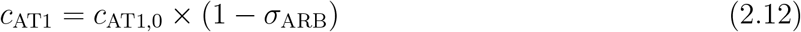

where *c*_AT1,0_ is the baseline value for *c*_AT1_ and *σ*_ARB_ is the percent inhibition of AT1R binding by the ARB. Similarly, to simulate treatment with an ACE inhibitor (ACEi), we let

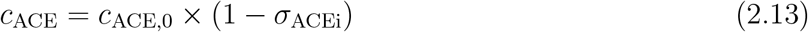

where *c*_ACE,0_ is the baseline value for *c*_ACE_ and *σ*_ACEi_ is the percent inhibition of ACE by the ACEi.

### 2.2 Calcium homeostasis and bone biology model

The mathematical model developed by Peterson and Riggs [13] consists of detailed bone, renal, and blood compartments, and predicts levels of PTH, calcitriol, bone cell activity, calcium, and phosphate. In this study, we primarily used the original model from Peterson and Riggs [13] for the calcium homeostasis and bone remodeling components with the following modifications from published model extensions:

- Gaweda et al. [16] extended this model to include other known regulatory mechanisms on calcium and phosphate homeostasis. In this study, we also included the impact of phosphate levels on parathyroid gland capacity as added by Gaweda et al. [16] (equation for *H*_5,11_ in Ref. [16]). Note that Gaweda et al. [16] used the Peterson and Riggs [13] model as is with their extensions so all other parameters and biological components are the same as in Ref. [13].
- A compartment for bone mineral density was added by Riggs et al. [14] and is included in the model in this study.
- Riggs et al. [14] added the impact of estrogen on bone remodeling and calcium homeostasis to the model in Ref. [13]. The authors included the impact of estrogen on latent and active transforming growth factor-*β* (TGF-*β*), osteoblast signaling, and renal calcium handling. TGF-*β* is a key cytokine in bone remodeling that regulates osteoblast and osteoclast proliferation and activity. Estrogen inhibits latent TGF-*β* formation, while stimulating its activation. TGF-*β* impacts bone remodeling activity activity through stimulating the expression of RANK on osteoclasts, stimulating osteoclast apoptosis, stimulating pre-osteoblast formation, and inhibiting osteoblast proliferation. Estrogen directly inhibits both the formation of pre-osteoblasts as well as the apoptosis rate of osteoblasts. Riggs et al. [14] also included the impact of estrogen on renal calcium reabsorption by decreasing renal calcium reabsorption with declining estrogen levels post-menopause. All of these estrogen impacts were included in the model use in this study.

A model schematic is shown in Fig. 2.1. The model equations are listed in Appendix D.

### 2.3 Composite calcium homeostasis, bone remodeling, and RAS model

To formulate a composite model, we connected the estrogen-RAS model (Section 2.1) with the calcium homeostasis and bone biology model (Section 2.2) through known regulatory mechanisms. Specifically, the composite model includes the impact of AT1R-bound Ang II on PTH secretion (Section 2.3.1), the impacts of extracellular calcium levels and the calciotropic hormones on renin secretion (Section 2.3.2), and the impact of AT1R-bound Ang II on bone remodeling (Section 2.3.3).

To model those interactions, we follow the approach inRef. [13] and characterize the changes in responses influenced by a given stimulant using hyperbolic functions:

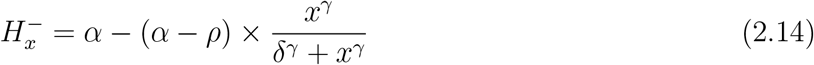

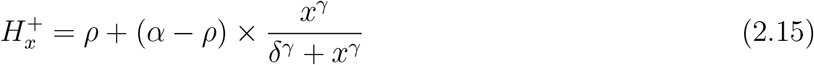

where *x* denotes the stimulus variable, *α* is the maximum anticipated response, *γ* is the steepness of the response (note when *γ* = 1 the functions are sigmoidal), *δ* is the value of *x* that produces the half-maximal response, and *ρ* is the minimum anticipated response. The minus or plus superscript in *H* denotes whether the response is a decrease or increase from the baseline steady state, respectively. Note that when *ρ* = 0, the response is a classical *E*_max_ response often used in modeling in pharmacology (see Ref. [36]). Parameter values for the equations added in this study are listed in Table 2.2. Additional parameter values can be found in Refs. [13, 14, 16] or using our publicly available model code (see Section 2.5).

**Table 2.2:**
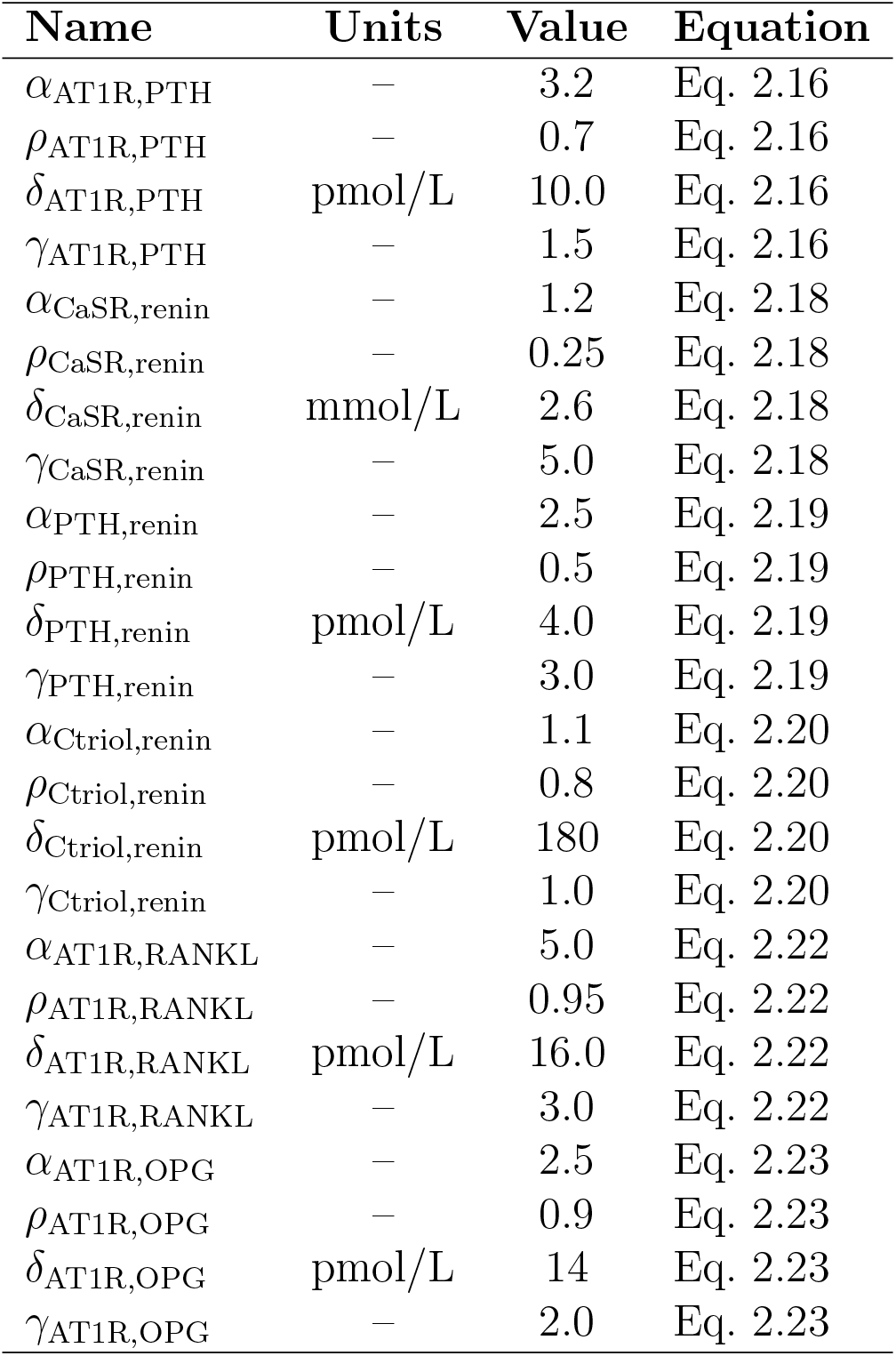
Parameter values for coupling the RAS components to the calcium homeostasis and bone remodeling model. Additional parameters can be found in Refs. [13, 16].

| Name | Units | Value | Equation |
| --- | --- | --- | --- |
| $\alpha_{\text{AT1R,PTH}}$ | — | 3.2 | Eq. 2.16 |
| $\rho_{\text{AT1R,PTH}}$ | — | 0.7 | Eq. 2.16 |
| $\delta_{\text{AT1R,PTH}}$ | pmol/L | 10.0 | Eq. 2.16 |
| $\gamma_{\text{AT1R,PTH}}$ | — | 1.5 | Eq. 2.16 |
| $\alpha_{\text{CaSR,renin}}$ | — | 1.2 | Eq. 2.18 |
| $\rho_{\text{CaSR,renin}}$ | — | 0.25 | Eq. 2.18 |
| $\delta_{\text{CaSR,renin}}$ | mmol/L | 2.6 | Eq. 2.18 |
| $\gamma_{\text{CaSR,renin}}$ | — | 5.0 | Eq. 2.18 |
| $\alpha_{\text{PTH,renin}}$ | — | 2.5 | Eq. 2.19 |
| $\rho_{\text{PTH,renin}}$ | — | 0.5 | Eq. 2.19 |
| $\delta_{\text{PTH,renin}}$ | pmol/L | 4.0 | Eq. 2.19 |
| $\gamma_{\text{PTH,renin}}$ | — | 3.0 | Eq. 2.19 |
| $\alpha_{\text{Ctriol,renin}}$ | — | 1.1 | Eq. 2.20 |
| $\rho_{\text{Ctriol,renin}}$ | — | 0.8 | Eq. 2.20 |
| $\delta_{\text{Ctriol,renin}}$ | pmol/L | 180 | Eq. 2.20 |
| $\gamma_{\text{Ctriol,renin}}$ | — | 1.0 | Eq. 2.20 |
| $\alpha_{\text{AT1R,RANKL}}$ | — | 5.0 | Eq. 2.22 |
| $\rho_{\text{AT1R,RANKL}}$ | — | 0.95 | Eq. 2.22 |
| $\delta_{\text{AT1R,RANKL}}$ | pmol/L | 16.0 | Eq. 2.22 |
| $\gamma_{\text{AT1R,RANKL}}$ | — | 3.0 | Eq. 2.22 |
| $\alpha_{\text{AT1R,OPG}}$ | — | 2.5 | Eq. 2.23 |
| $\rho_{\text{AT1R,OPG}}$ | — | 0.9 | Eq. 2.23 |
| $\delta_{\text{AT1R,OPG}}$ | pmol/L | 14 | Eq. 2.23 |
| $\gamma_{\text{AT1R,OPG}}$ | — | 2.0 | Eq. 2.23 |

#### 2.3.1 AT1R-bound Ang II impact on PTH secretion

AT1R-bound Ang II concentration, denoted by [AT1R-AngII], increases PTH secretion from the parathyroid gland [6]. To model this effect, we let 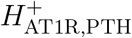 denote the impact of AT1R-bound Ang II on PTH secretion:

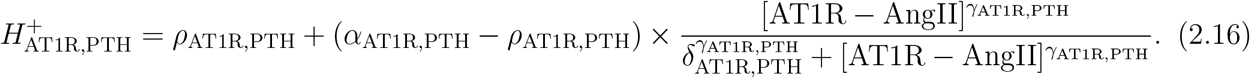

where the parameters correspond to the function form of type Eq. 2.15. PTH concentration ([PTH]) in the blood is modeled by

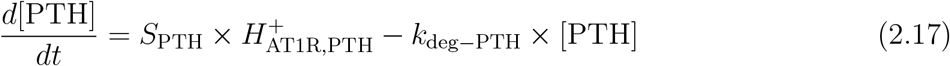

where *S*_PTH_ is the secretion of PTH from the parathyroid gland and *k*_deg*−*PTH_ is degradation as in Ref. [13]. The parameters for Eq. 2.16 were fit using the Ang II only experiment data from Grant et al. [37] (Section 2.3.4). The parameter values are listed in Table 2.2.

#### 2.3.2 Calciotropic hormones, calcium, and renin secretion

Renin is secreted from specialized cells in the kidneys called juxtaglomerular cells. Renin secretion, denoted by *R*_sec_, is impacted by [PTH], calcitriol ([Ctriol]), and extracellular calcium levels. Specifically, increased extracellular calcium and [Ctriol] inhibit renin secretion, while [PTH] stimulates renin secretion [6].

Extracellular calcium levels are detected by the calcium sensing receptor (CaSR) in the juxtaglomerular cells [38]. To model the impact of CaSR on *R*_sec_ we let:

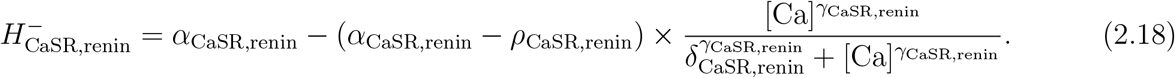

Ortiz-Capisano et al. [39,40] found that renin release was about 40% lower when using a calcimimetic or at elevated calcium levels in the juxtaglomerular cells of the kidneys. We used this result to estimate parameters so that at high calcium levels, renin release is decreased by about 40%.

To model stimulation of *R*_sec_ by [PTH] we let:

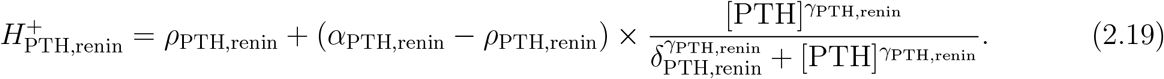

The parameters (listed in Table 2.2) were fit using the PTH only infusion experiment from Grant et al. [37] (Section 2.3.4).

To model inhibition of *R*_sec_ by calcitriol we let:

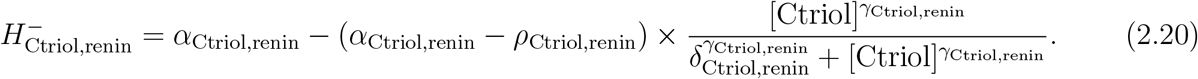

The parameters were chosen (listed in Table 2.2) based on the data from Tomaschitz et al. [41] that related calcitriol levels with renin concentration. Taken together, we incorporate the combined impact of calcium, PTH, and calcitriol on renin secretion by modifying Eq. 2.4 to

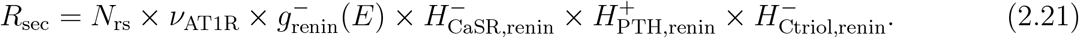

#### 2.3.3 AT1R-bound Ang II activation of RANKL and OPG

Bone remodeling is primarily regulated by osteoblasts and osteoclasts. Osteoblasts develop the bone matrix, while osteoclasts break down bone tissue. RANKL (receptor activator of nuclear factor kappa-B ligand) is expressed by osteoblasts and binds to its receptor RANK (receptor activator of nuclear factor kappa-B) on osteoclasts to promote osteoclast differentiation and activation. Osteoprotegerin (OPG) is a decoy receptor that binds to RANKL, preventing its binding with RANK and thereby inhibiting osteoclast activity. The balance between RANKL and OPG is essential for maintaining bone homeostasis as an increase in RANKL or decrease in OPG can lead to excessive or reduced bone resorption, respectively. This system is often referred to as the RANK-RANKL-OPG system and is illustrated in Fig. 2.1.

Shimizu et al. [7] reported that in ovariectomized rats, infusion of Ang II increased RANKL expression and led to a significant decrease in bone volume, but this impact was abolished by treatment with an AT1R blocker and not an AT2R blocker. Their results showed that RANKL production is activated by AT1R-bound Ang II levels. Additionally, Shimizu et al. [7] found that OPG expression is also increased in osteoblast cells due to infusion of Ang II, but to a much lesser extent than RANKL.

The impact of AT1R-bound Ang II on RANKL and OPG expression is modeled using the form in Eq. 2.15 for activation so that:

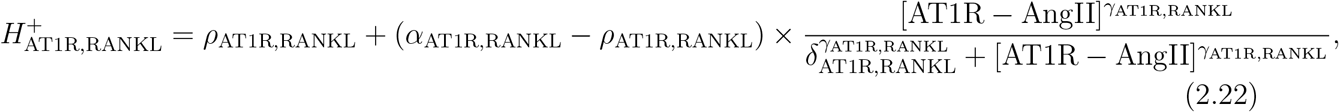

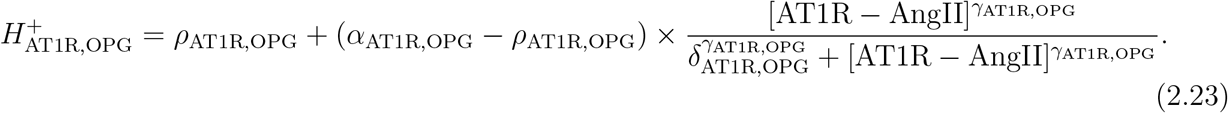

These are included in the equations for the production of RANKL and OPG, respectively (Eqs. D.38 and D.40 in Appendix D). The values for *α*_AT1R,RANKL_ and *α*_AT1R,OPG_ represent the highest possible amount of activation of RANKL and OPG by AT1R binding, respectively. These values were set to the fold-change in RANKL and OPG expression after Ang II infusion in Shimizu et al. [7]. The parameter values are listed in Table 2.2.

#### 2.3.4 Parameter fittingt o P TH and Ang II infusion experiments

Grant et al. [37] conducted infusion experiments in healthy volunteers to determine the relationship between the RAS and PTH in humans. To test this, the authors gave healthy volunteers 1 hour infusions on different days of (i) Ang II only, (ii) PTH (i.e., teriparatide) only, and (iii) Ang II and PTH together. We simulated these experiments and used the data from this study to determine parameters values (listed in Table 2.2) for Eqs. 2.16, 2.18, and 2.19.

In their experiments, Ang II infusions were given in graded doses of 1, 3, and 10 ng/kg*·*min for consecutive infusions of 20 minutes each. We simulated Ang II infusion by letting *k*_inf_ = 1 ng/kg*·*min (Eq. 2.9) for the first 20 minutes, 3 ng/kg·min for the next 20 minutes, and 10 ng/kg·min for the final 20 minutes of the 1 hour infusion to simulate this infusion type. For PTH infusions we used the pharmacokinetic parameters from Peterson and Riggs [13] with a dose of 200 U of teriaparatide as reported by Grant et al. [37]. To fit the parameters, we used the fmincon function from MATLAB using the interior-point algorithm to minimize the difference between the predicted values and the data for PRA, intact PTH, and [Ang II] reported in Grant et al. [37]. In all of these simulations we let *E*=1 so that there are no estrogen effects.

### 2.4 Sobol sensitivity analysis

Mathematical models of complex biological systems, like the one used in this study, involve a large number of parameters with unknown or uncertain values. To investigate the impact of variations in parameter values on model output, we can use methods such as sensitivity analysis [42, 43]. Global sensitivity analysis methods evaluate the model at many points in the parameter space, providing an assessment of how individual parameters contribute to variations in model output across the parameter space, including interactions with other parameter changes [42–44].

The Sobol method is a variance-based global sensitivity analysis technique that decomposes the variance in the model output into contributions from each individual parameter as well as its interactions with other parameters [42, 45]. The results of a Sobol sensitivity analysis are called “Sobol sensitivity indices” (or “Sobol indices” for short) and are computed for each parameter included in the analysis. First-order Sobol indices (denoted by *S*_first_) are the fraction of the variance in the model output the individual parameter contributes to. Total order Sobol indices (denoted by *S*_total_) quantify the fraction of output variance the parameter contributes to, including interactions with other parameter changes. By evaluating Sobol indices, we can identify which parameters are the most influential on model output. For more information about Sobol sensitivity analysis and its use in applications in biology, we refer readers to Refs. [42–45].

We conducted a Sobol sensitivity analysis on the full composite model (described in Section 2.3). The model outputs we considered are bone mineral density, AT1R-bound Ang II, PTH, calcitriol, extracellular calcium, and renin concentration over a simulation time from 20 to 80 years with agerelated estrogen decline (i.e., same as the “no RASi” simulations in Figs. 3.2, 3.3). We computed Sobol indices at every 10 years of simulation time. The full composite model involves 181 parameters, so running a Sobol sensitivity analysis over all the parameters can lead to large numerical errors and computation time. To reduce the number of parameters used in the full Sobol sensitivity analysis, we applied a two-step approach:

1. We first conducted a Sobol sensitivity analysis varying all parameters only in the calcium homeostasis and bone biology model described in Section 2.2 by ±10%. This includes the baseline biological components of bone remodeling (including the detailed impacts of RANKL, RANK, OPG, and TGF*β* on osteoblast and osteoclast activity), PTH secretion and degradation, calcitriol synthesis and degradation, renal calcium and phosphate reabsorption, intestinal calcium and phosphate absorption, and extracellular calcium and phosphate levels. We screened the 121 parameters included in this analysis for parameters that had an *S*_first_ value of greater than 1% in at least one of the outputs of interest (bone mineral density, AT1R-bound Ang II, PTH, calcitriol, or calcium) at some point in the simulation. This resulted in 30 sensitive parameters.
2. We then included all the new parameters from this study (listed in Tables 2.1 and 2.2) with the 30 parameters selected from the initial screening in Step 1, resulting in 59 total parameters. We conducted a Sobol sensitivity analysis for the full composite model with parameters varied by *±*25% of their baseline value.

The results of the Sobol sensitivity analysis are described in Section 3.3.

### 2.5 Code availability

All model code used in this study is available at https://github.com/Layton-Lab/est-ras-bone. Model simulations and code were conducted using MATLAB R2022a. The Sobol sensitivity analysis was conducted using Simbiology. Data originally presented in graphs from published papers (e.g., the data from Grant et al. [37] shown in Fig. 3.1) were extracted using Web Plot Digitizer (version 4.7).

**Figure 3.1:**
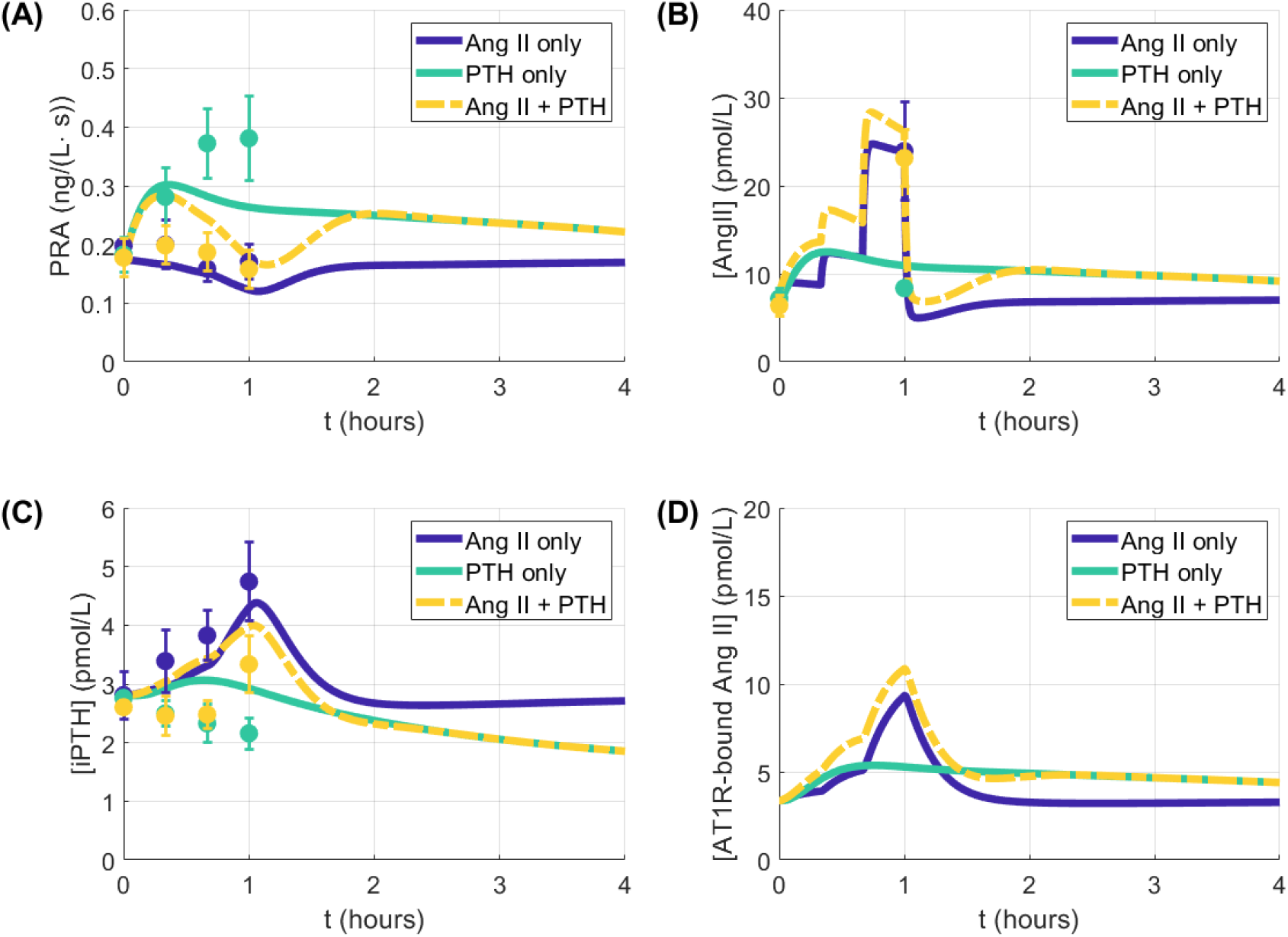
Simulation results for infusion of Ang II (“Ang II only”), PTH (“PTH only”), and Ang II and PTH together (“Ang II + PTH”). Results for (A) plasma renin activity, (B) angiotensin II plasma concentration, (C), intact parathyroid hormone, and (D) angiotensin receptor type 1-bound Ang II are shown for during and after 1 hour infusions. Lines indicate simulations, while the dots and error bars in panels A–C are from the relevant experimental data presented in Grant et al. [37]. Ang II: angiotensin II, AT1R: angiotensin receptor type 1, iPTH: intact parathyroid hormone, PRA: plasma renin activity

## 3 Results

### 3.1 PTH and Ang II infusion experiments simulations

We conducted simulations of infusion of (i) Ang II, (ii) PTH, and (iii) PTH and Ang II together (described in Section 2.3.4). Model results, together with the experimental data reported for the clinical studies done by Grant et al. [37], are shown in Fig. 3.1.

When Ang II is infused in isolation, Ang II levels rise resulting in an increase in AT1R-bound Ang II levels (Fig. 3.1B, D), which stimulates PTH secretion so that endogenous PTH (intact PTH, denoted by [iPTH]) increases (Fig. 3.1C). Grant et al. [37] reported that during an Ang II infusion, [iPTH] increased, while PRA decreased slightly likely due to the negative feedback on renin secretion by AT1R-bound Ang II (Fig. 3.1 “Ang II only” data). Model simulations capture both the rise in [iPTH] (Fig. 3.1C “Ang II only”) and slight decrease in PRA (Fig. 3.1A “Ang II only”) during infusion.

When infusing PTH in healthy volunteers, PRA nearly doubled on average (Fig. 3.1A “PTH only” data) [37]. Model simulations predicted a 69% increase in PRA (Fig. 3.1A “PTH only”). While total PTH levels increased substantially during the PTH infusion experiment, endogenous PTH production steadily decreased (Fig. 3.1C “PTH only” data). Model simulations did not result in a significant decrease [iPTH]. That discrepancy may be attributed to a more detailed impact of PTH on renal calcium reabsorption, that is not represented in the model, which would increase calcium levels and thus decrease PTH secretion via the CaSR [46].

When infusing both Ang II and PTH together in healthy volunteers, the authors reported that Ang II levels were similar to those reported in the Ang II only experiment, likely due to PTH having only a small impact on Ang II levels [37]. In our model simulations, Ang II levels were slightly higher in the Ang II and PTH infusion experiment than the Ang II only experiment, but were still within the range reported by Grant et al. [37] at the end of the 1 hour infusion time (Fig. 3.1B “Ang II + PTH”). Grant et al. [37] reported that PRA levels remained near baseline for the Ang II and PTH infusion experiment. In our model simulations, the “Ang II + PTH” simulations resulted in a transient increase in PRA but declines to near the levels prior to infusion by the end of the simulation time (Fig. 3.1A).

To summarize, model parameters were chosen to fit the observed responses of key RAS components and PTH following Ang II and PTH infusions with baseline estrogen levels.

### 3.2 Impact of estrogen decline and RAS inhibitors

How do declining estrogen levels in post-menopausal women impact the RAS? How do changes in the RAS and declining estrogen levels contribute to altered bone remodeling? Does bone mineral density improve with RAS inhibition? To investigate these questions, we conducted simulations of the composite calcium homeostasis, bone remodeling, and estrogen-RAS model with estrogen decline observed in post-menopausal women with and without drug interventions.

In each simulation, estrogen decline starts at 50 years, which is about the average age of menopause onset in women. We conducted simulations with key RAS inhibitors (ACEi and ARB) starting at age 60 years, unless otherwise indicated. For ARB simulations, we let *σ*_ARB_ (Eq. 2.12) vary from 88–98.3% as reported for different types and typical dose levels of ARBs by Hallow et al. [18]. For ACEi, we conducted simulations with *σ*_ACEi_ (Eq. 2.13) varying from 94–97.2% as reported for common types of ACEi and doses by Hallow et al. [18]. The results are shown in Figs. 3.2 and 3.3.

**Figure 3.2:**
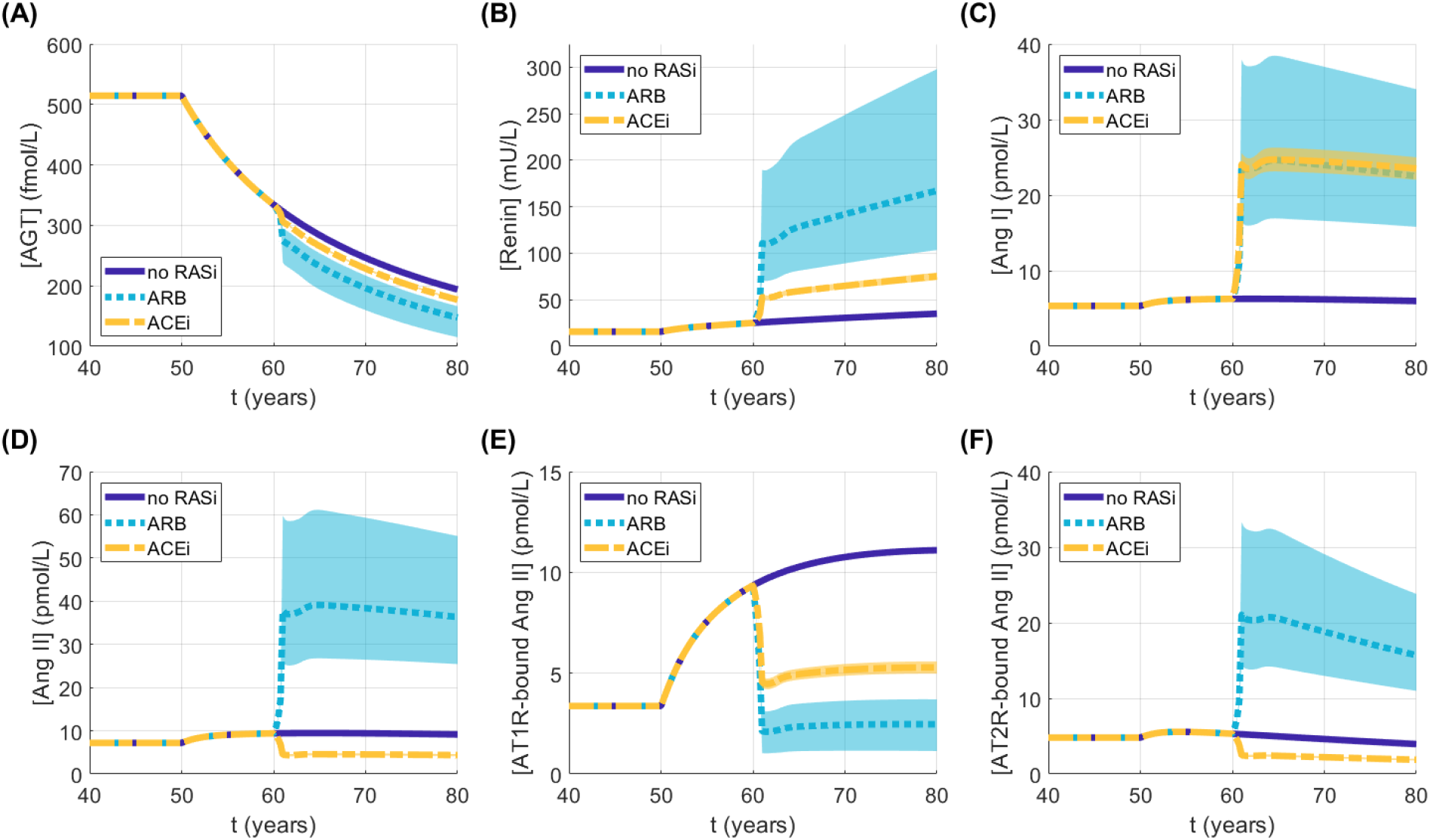
Impact of post-menopausal estrogen decline and renin-angiotensin system inhibitors on the renin-angiotensin system. For the angiotensin receptor blocker simulations (labeled “ARB” in the legend), σ_ARB_ (Eq. 2.12) was varied between 88–98.3% based on the ranges reported in Ref. [18]. For the angiotensin-converting enzyme inhibitor simulations (labeled “ACEi” in the legend), σ_ACEi_ (Eq. 2.13) was varied between 94–97.2% as reported in Ref. [18]. Lines represent the mean values over the varied σ_i_ values for the respective treatment with shaded area representing the range. AGT: angiotensinogen, Ang I: angiotensin I, Ang II: angiotensin II, AT1R: angiotensin type 1 receptor, AT2R: angiotensin type 2 receptor, RAS: renin angiotensin system, RASi: RAS inhibitor, ARB: angiotensin receptor blocker, ACEi: angiotensin-converting enzyme inhibitor

ACEi and ARB inhibit ACE activity and AT1R binding, respectively. While these are different components of the RAS, both drugs result in reduced levels of AT1R-bound Ang II (Fig. 3.2E). The impact of these drugs on the various components of the RAS are drug and dose-dependent. The varied range in values of *σ*_ACEi_ had a negligible effect on our model simulation results (Fig. 3.2, “ACEi”). In the ARB simulations, the impact on the RAS has a wider variation based on the value of *σ*_ARB_ (Fig. 3.2, “ARB”). Note that we simulated a wider range for *σ*_ARB_ than *σ*_ACEi_.

We first consider the impact of ACEi and ARB on the RAS components. ARB directly inhibit the binding of Ang II to AT1R. As a result, AT1R-bound Ang II levels decreased by about 66–89%, depending on efficacy (Fig. 3.2E, “ARB” vs. “no RASi”). Recall that AT1R-bound Ang II inhibits renin secretion (Fig. 2.1). Hence, decreased AT1R-bound Ang II levels result in increased renin levels (Fig. 3.2B) as well as increased downstream Ang I (Fig. 3.2C) and Ang II levels (Fig. 3.2D). The lower AT1R binding along with increased Ang II levels results in more Ang II binding to AT2R, so that AT2R-bound Ang II levels increase by about 4-fold on average (Fig. 3.2F “ARB” vs. “no RASi”‘). ACEi impact the RAS by inhibiting the activity of ACE, which is the enzyme that converts Ang I into Ang II (Fig. 2.1). Ang II levels decreased by an average of 52% in the ACEi simulations despite increased Ang I levels (Fig. 3.2D, C “ACEi” vs. “no RASi”). As a result, both AT2R-bound Ang II and AT1R-bound Ang II decreased by about 52% from the no RAS inhibitor case (Fig. 3.2E–F, “ACEi” vs. “no RASi”).

Next, we analyze the age-related estrogen decline effect (Appendix A) on bone mineral density and its response to ACEi and ARB. Estrogen maintains bone density by regulating factors involved in bone remodeling that promote the differentiation and activation of osteoblasts, while inhibiting osteoclast activity (Fig. 2.1). When estrogen levels decline, osteoclast activity increases, which increases bone resorption, ultimately resulting in a decrease in bone mineral density (Fig. 3.3A–B “no RASi”). The bone mineral density simulations results with the decline in estrogen levels is consistent with the data reported for aging women in Looker et al. [47] (Fig. 3.3A “no RASi” data). This decrease is a result of a combination of the direct impacts of estrogen on bone remodeling and increased AT1R-bound Ang II levels, which stimulates RANKL production to bind with RANK (Fig. 3.3C; Fig. 3.2E “no RASi”). RANK-RANKL stimulates osteoclast differentiation and activation. Throughout the estrogen decline simulations, the calciotropic hormones, PTH and calcitriol, as well as key electrolytes, calcium and phosphate, remained within normal range (Fig. B.2).

Age-related decreases in estrogen levels result in 4.5-fold increase in osteoclast activity (Fig. 3.3B “no RASi”). However, with ARB, AT1R-bound Ang II levels decrease (Fig. 3.2E), resulting in reduced RANKL production, so that osteoclast levels return to about 40% above baseline levels (Fig. 3.3B,C). As a result, by the end of the simulation time (i.e, *t* = 80 years), bone mineral density is at about 90% of baseline for ARB users versus 75% before estrogen decline with no treatment (Fig. 3.3A “ARB” vs. “no RASi”). ACEi also decreased AT1R-bound Ang II levels by a lesser amount (Fig. 3.2E) so that osteoclast levels are lowered to about 2-fold baseline levels. This results in a slight improvement in bone mineral density to about 85% of the level before estrogen decline (Fig. 3.3A “ACEi”). We observed that the range in the amount of AT1R-bound Ang II inhibition due to varied ARB efficacy did not have a large impact the change in bone levels on these drugs (Fig. 3.3; Fig. 3.2). Thus, the improvement to bone mineral density may not depend significantly on the dose of the RAS inhibitors.

**Figure 3.3.**
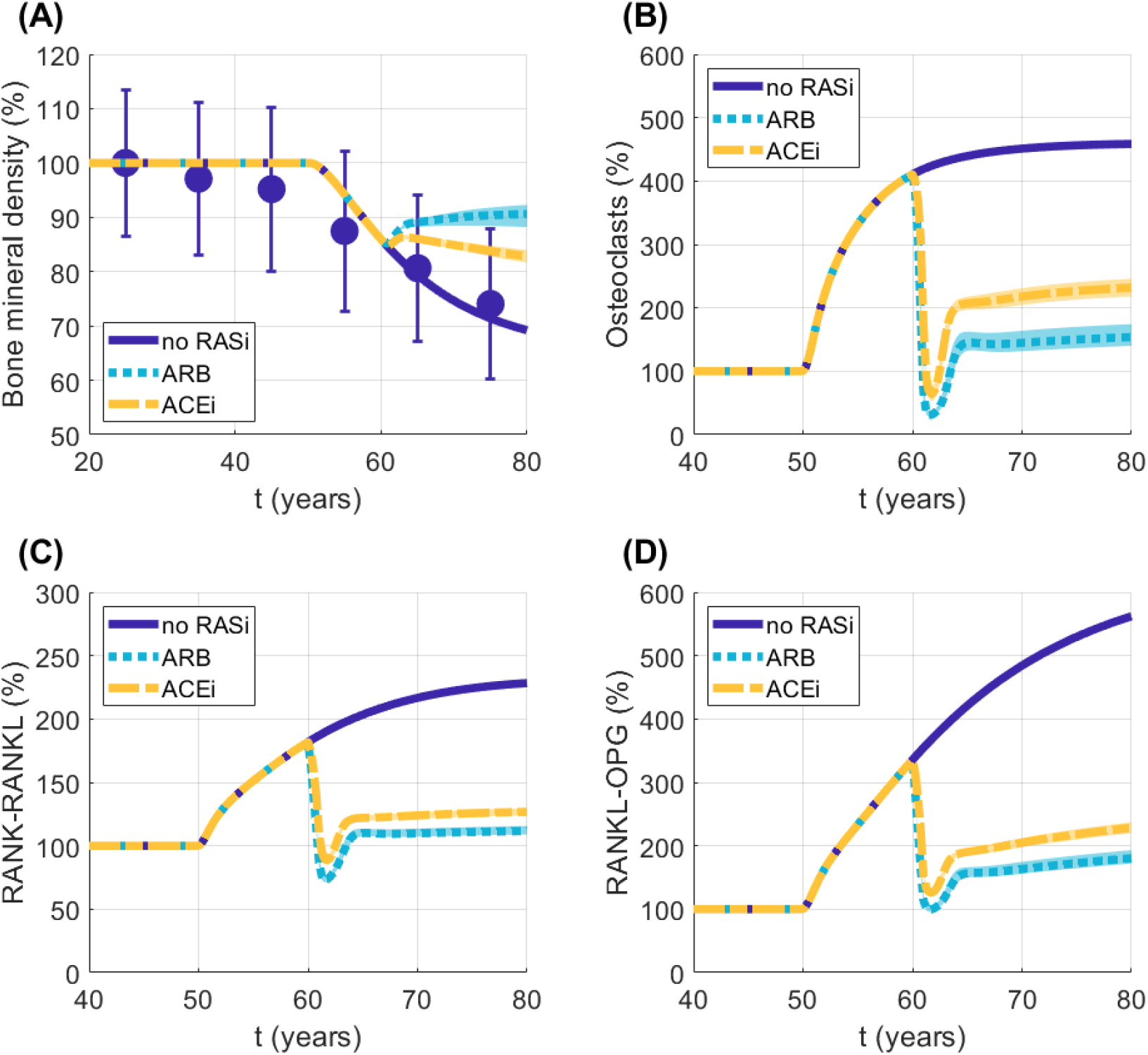
Impact of post-menopausal estrogen decline and RAS inhibitors on (A) bone mineral density, (B) osteoclast levels, (C) RANK-RANKL levels, and (D) RANK-OPG levels. The values are normalized to their baseline levels and shown as a percent of the baseline value. For the angiotensin receptor blocker simulations (labeled “ARB” in the legend), σ_ARB_ (Eq. 2.12) was varied between 88–98.3% based on the ranges reported in Ref. [18]. For the angiotensin-converting enzyme inhibitor simulations (labeled “ACEi” in the legend), σ_ACEi_ (Eq. 2.13) was varied between 94–97.2% as reported in Ref. [18]. Lines represent the mean values over the varied σ_i_ values for the respective treatment with shaded area representing the range. Data in panel (A) are used from the female data for femoral neck bone mineral density in Looker et al. [47]. The simulations and data are for femoral neck bone mineral density. BMD: bone mineral density, RANK: receptor activator of NF-kappaB, RANKL: RANK ligand, OPG: osteoprotegerin, RASi: RAS inhibitor, ARB: angiotensin receptor blocker, ACEi: angiotensin-converting enzyme inhibitor

We conducted simulations of RAS inhibitor treatment starting at different ages with age-related estrogen decline as in Eq. 2.1. In these simulations, we let *σ*_ARB_ = 93% and *σ*_ACEi_ = 95% based on the average value for the respective drugs from Ref. [18]. We did this because varying *σ*_*i*_ did not have much impact on bone mineral density (Fig. 3.3A). The results are shown in Fig. 3.4. In all simulations, bone mineral density improved on both the ARB and ACEi drugs with a greater improvement with ARB treatment. When the administration of ARB or ACEi was started before the onset of estrogen decline (i.e., starting a RAS inhibitor before 50 years old), bone mineral density increased from baseline by about 5% (Fig. 3.4A1–A2). When estrogen decline started at 50 years, the bone mineral density decreased, but stays above 95% for patients on ARB (Fig. 3.4A1). When the administration of ACEi began before estrogen decline, bone mineral density declined below 90% of baseline during age-related estrogen decline (Fig. 3.4A2). Our model simulations suggest that early RAS inhibition prevents bone loss driven by age-related estrogen decline.

**Figure 3.4:**
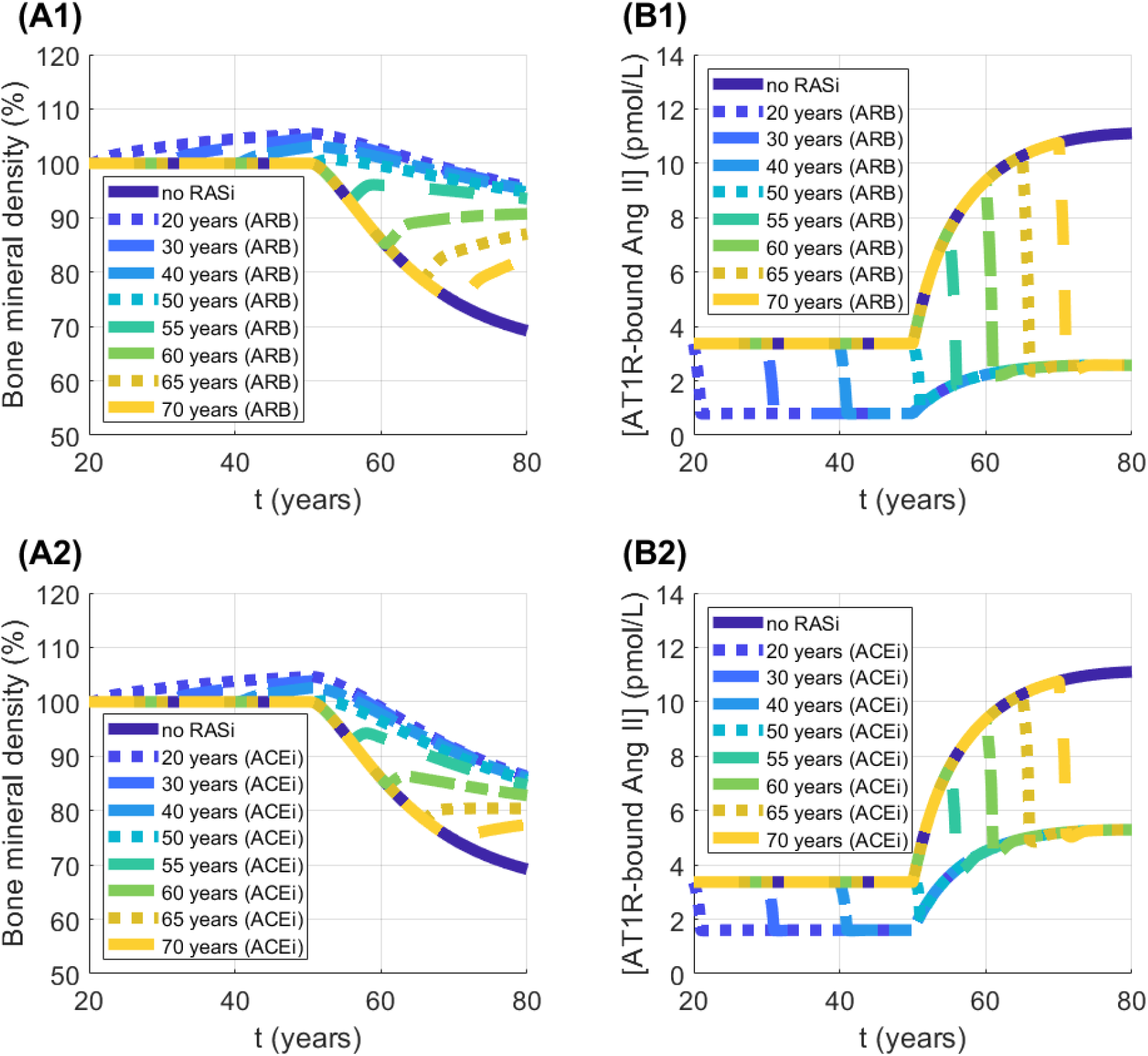
Model simulation results for varied starting age of angiotensin receptor blocker (A1–B1) and angiotensin-converting enzyme inhibitor (A2–B2). σ_ARB_ = 93% in all simulations for ARB and σ_ACEi_ = 95.6% in all ACEi simulations. The bone mineral density is femoral neck bone mineral density. AT1R: angiotensin type 1 receptor, ARB: angiotensin receptor blocker, ACEi: angiotensin-converting enzyme inhibitor, RASi: RAS inhibitor

In summary, the RAS, calcium, and bone remodeling model developed in this study represents the expected decrease in bone mineral density due to age-related estrogen decline. This is driven by both the direct impacts of estrogen on bone remodeling and calcium regulation, as well as RAS activation by estrogen decline. Model simulations suggest that ARBs offer stronger protection for bone health than ACEi, and that result is consistent when starting treatment at different ages.

### 3.3 Sensitivity analysis

We conducted a Sobol sensitivity analysis for the full composite model during age-related estrogen decline in women (see Section 2.4 for details). The parameters with the largest average *S*_total_ values in Fig. 3.5A are the parameters that have the largest impact on variation in bone mineral density. The top five parameters, based on the average *S*_total_ value, are all related to the regulation of RANK and RANKL (Fig. 3.5A). The RANK-RANKL complex directly impacts bone remodeling the stimulating osteoclasts (Fig. 2.1). Specifically, the parameters *k*_in,L_ and *k*_in,RNK_ are baseline production rates of RANKL and RANK, respectively. The parameter *E*_max,L*−*PTH_ relates to the impact of PTH on RANKL production. The parameters *k*_3_ and *k*_4_ are the binding and unbinding rate of the RANK-RANKL complex, respectively. The next parameter with the largest impact on bone mineral density is *ρ*_AT1R,L_, which represents the minimum impact of AT1R on RANKL production (Eq. 2.22). *δ*_CaSR,renin_ represents the sensitivity of renin secretion to calcium levels. This is the parameter with the largest impact on [AT1R-bound Ang II] (Fig. 3.5B). In summary, the top parameters that impact bone mineral density all primarily relate to RANK, RANKL and RANK-RANKL binding.

**Figure 3.5:**
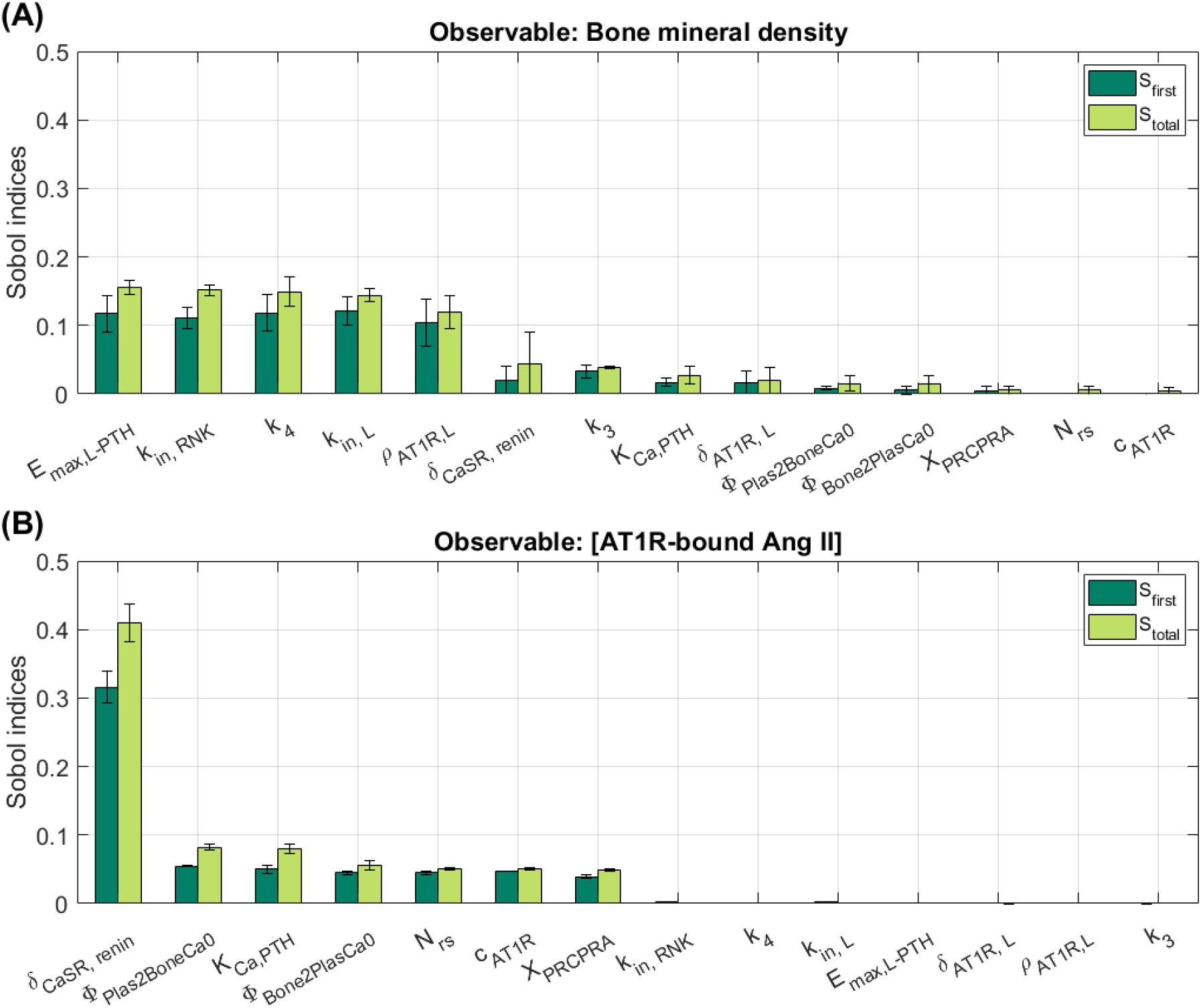
Mean (bar) and standard deviation (error bar) of first and total order Sobol indices (denoted by S_first_ and S_total_ in legend, respectively) of full model ran from age 20 to 80 years with estrogen decline for output variables (A) femoral neck bone mineral density and (B) AT1R-bound Ang II. Parameters shown had a maximal value of S_total_ > 0.05 for at least one output variable. A description of the parameters is given in Table C.1. AT1R: angiotensin type 1 receptor, Ang II: angiotensin II

AT1R-bound Ang II is the key biological component that connects the RAS with the calcium homeostasis and bone remodeling model (Fig. 2.1). The parameters with the 7 largest average *S*_first_ and *S*_total_ values for the model output of [AT1R-bound Ang II] are *δ*_CaSR,renin_, Φ_Plas2BoneCa0_, *K*_Ca,PTH_, Φ_Bone2PlasCa0_, *N*_rs_, *c*_AT1R_, and *X*_PRCPRA_ (Fig. 3.5B). *δ*_CaSR,renin_ represents half-maximal response of renin secretion to calcium levels. Φ_Plas2BoneCa0_ and Φ_Bone2PlasCa0_ represent the baseline fluxes of calcium between the plasma and bone and both impact variation in calcium and renin levels (Fig. 3.6). Calcium level changes affect the RAS via the CaSR in the renin-secreting juxtaglomerular cells of the kidneys. *K*_Ca,PTH_ is the half-maximal response of PTH secretion to calcium levels as mediated by the CaSR. This parameter plays a key role in variations in model output for [PTH], [Calcitriol], [Calcium], and [Renin] (Fig. 3.6 and 3.7), which all regulate the RAS. *N*_rs_, *c*_AT1R_, and *X*_PRCPRA_ are all directly related to the RAS (Section 2.1.1). Specifically, *N*_rs_ determines baseline renin secretion, *c*_AT1R_ is the binding rate for AT1R, and *X*_PRCPRA_ is baseline conversion rate from renin concentration to activity. Our sensitivity analysis shows that variations in [AT1R-bound Ang II] are driven by the CaSR impact on renin secretion and the RAS cascade.

**Figure 3.6:**
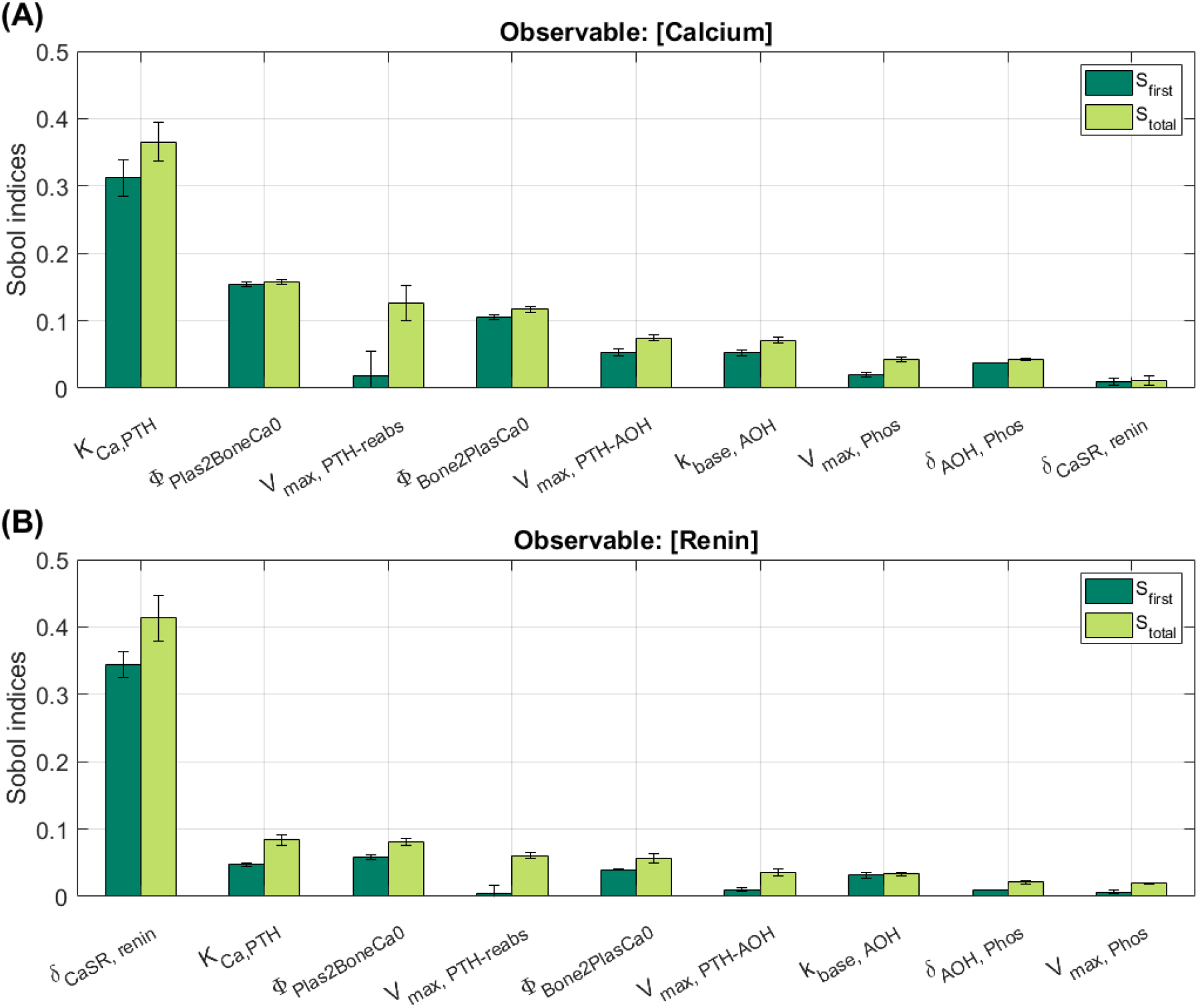
Mean (bar) and standard deviation (error bar) of first and total order Sobol indices (denoted by S_first_ and S_total_ in legend, respectively) of full model simulation from age 20 to 80 years with estrogen decline. Parameters shown had a maximal value of S_total_ > 0.05 for at least one output variable. A description of the parameters is given in Table C.1.

The calciotropic hormones are the primary regulators of calcium and bone remodeling. Additionally, these hormones have a bidirectional relationship with the RAS (Fig. 2.1). In our Sobol sensitivity analysis, the parameters with the largest impact on PTH had high *S*_total_ values relative to the respective *S*_first_ values, likely due to the large number of regulators on PTH; these parameters are *V*_max,PTH*−*reabs_, *K*_Ca,PTH_, and *V*_PTH*−*AOH_ (Fig. 3.7A). *K*_Ca,PTH_ denotes the half-maximal response of PTH secretion driven by calcium sensing via the CaSR in the parathyroid glands. *V*_max,PTH*−*reabs_ relates to the impact of PTH on calcium reabsorption in the kidneys. This parameter has a high *S*_total_ value for [PTH] as well as [Calcium] (Fig. 3.6 and 3.7), likely due to the interactions between renal calcium reabsorption, calcium levels, and PTH secretion. *V*_PTH*−*AOH_ represents the maximal impact of PTH on 1*α*-hydroxylase, the enzyme that synthesizes calcitriol. PTH and calcitriol regulate one another, as calcitriol inhibits PTH secretion and PTH stimulates calcitriol production (Fig. 2.1). Both PTH and calcitriol are sensitive to the parameter *k*_base,AOH_ which is the baseline production rate of 1*α*-hydroxylase, and to the parameters Φ_Bone2PlasCa0_ and Φ_Plas2BoneCa0_, which denote the baseline rate of calcium exchange between the bone and plasma and vice versa (Fig. 3.7). The latter is likely due to the sensitivity of PTH to calcium levels, which are impacted by these two parameters as well (Fig. 3.6A).

**Figure 3.7:**
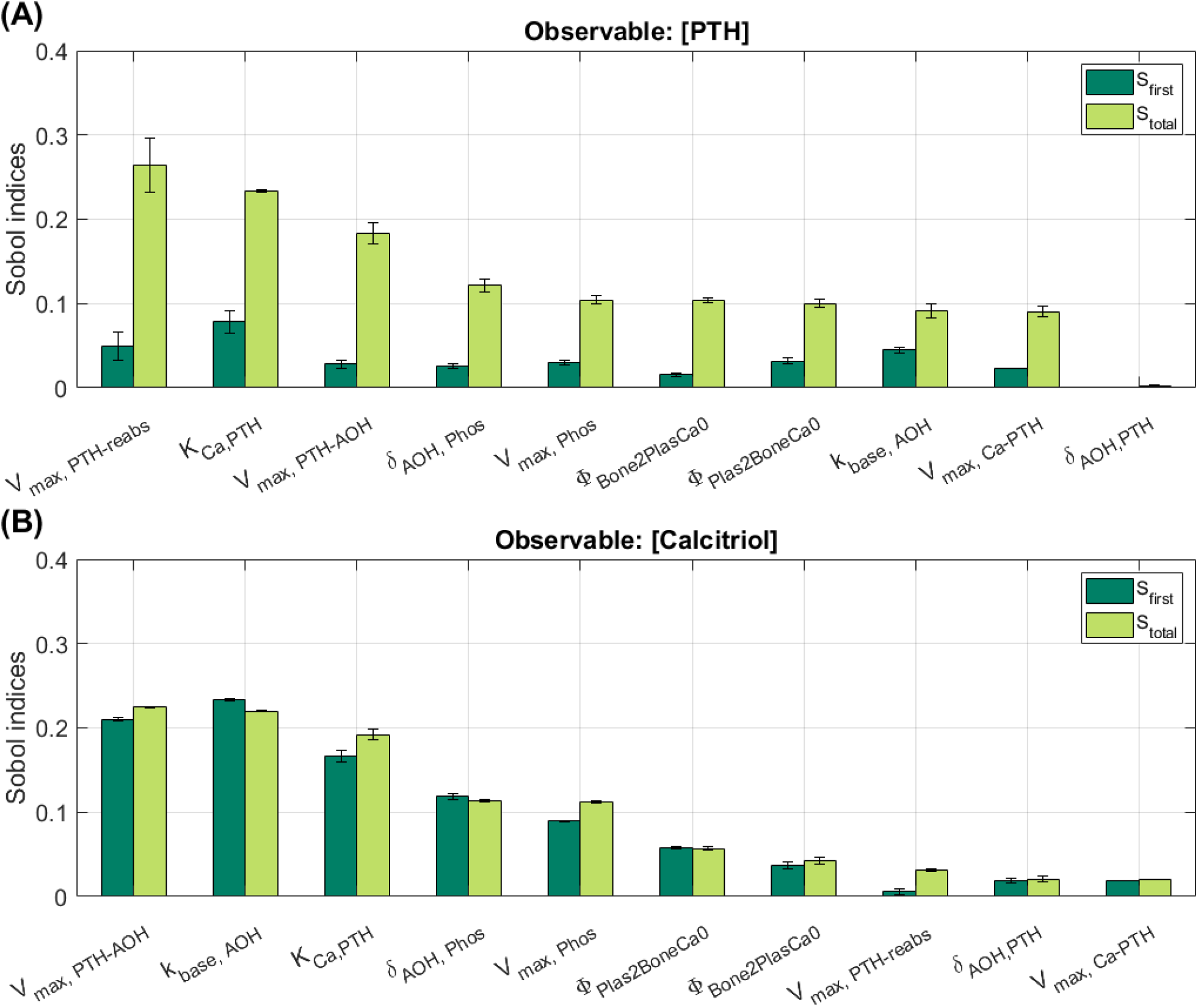
Mean (bar) and standard deviation (error bar) of first and total order Sobol indices (denoted by S_first_ and S_total_ in legend, respectively) of full model ran from age 20 to 80 years with estrogen decline for output variables (A) parathyroid hormone and (B) calcitriol. Parameters shown had a maximal value of S_total_ > 0.05 for at least one output variable. A description of the parameters is given in Table C.1. PTH: parathyroid hormone

The Sobol sensitivity analysis results shown here are the average (bar) and standard deviation (error bar) of *S*_first_ and *S*_total_ values over the full simulation time from age 20 to 80 years. We also analyzed the results over the simulation time from age 20 to 50, i.e., before estrogen decline. The parameters with the largest impact on model output were the same (results not shown) as those for the full simulation time. This suggests that the model output is most sensitive to parameters that represent the overall biological components of the composite model rather than parameters that represent the impact of estrogen on calcium regulation and bone remodeling.

In summary, our global sensitivity analysis shows that, in our model, bone remodeling is most sensitive to RANK-RANKL binding and that the RAS is sensitive to both calcium levels and the calciotropic hormones. These are the key biological components that impact RAS levels and bone remodeling.

## 4 Discussion

In this study we developed a mathematical model that includes the interconnected systems of estrogen, the RAS, PTH, calcitriol, calcium regulation, and bone remodeling. We used this model to:

- Investigate the impact of estrogen decline, as experienced in post-menopausal women, on the RAS, bone remodeling, and calcium homeostasis.
- Simulate potential therapeutic effects of ACEi and ARB on bone remodeling and calcium regulation.
- Assess the sensitivity of key model outputs to variations in the model’s many unknown parameters using a Sobol sensitivity analysis.

To the best of our knowledge, this is the first model to incorporate the direct effects of estrogen on the RAS, as well as the integrated effects of the RAS on calcium regulatory and bone remodeling systems. As such, our model development and analysis have brought major advances to the active area of using mathematical modeling to investigate bone remodeling and calcium regulation.

Given that the prevalence of hypertension increases markedly with aging, hypertension and osteoporosis often coexist. RAS inhibitors are widely used, safe, reasonably priced long term treatments for hypertension. RAS inhibitors have also been proposed as a treatment to improve bone density due to the relationship between the RAS and bone remodeling [10, 11]. Since RAS inhibitors are already widely used for the treatment of hypertension in aging populations, including post-menopausal women, it may be beneficial to consider which RAS inhibitor may prevent or improve bone loss.

We conducted model simulations to investigate the impacts of RAS inhibitors, specifically ACEi and ARB, on bone remodeling for various stages of age-related estrogen decline. Since treatment with an ARB results in a larger reduction in AT1R-bound Ang II for all dose levels, the improvement in bone mineral density was greater in simulations of ARB than ACEi (Fig. 3.2E, Fig. 3.3A). It seems that the targeted impact of ARB on AT1R binding results in a better improvement in bone mineral density during post-menopausal estrogen decline than ACEi, which inhibit Ang II production via ACE. These results were consistent when we varied the starting age for ACEi and ARB before and after estrogen decline (Fig. 3.4). In summary, our model simulations showed that reductions in AT1R-bound Ang II abundance results in reduced osteoclast activity, which prevents and improves declines in bone mineral density driven by low estrogen levels.

In a clinical study, Kim et al. [10] reported that patients that have used ARB only experienced fewer fractures than patients who have never used a RAS inhibitor. Similarly, Solomon et al. [48] reported reduced fracture risk in users of ARB, while ACEi users had no change in fracture risk. The systemic review by Wu et al. [12] reported that lower fracture risk is consistently reported in users of ARB, but not ACEi. In contrast, our mathematical model predicted that bone mineral density is increased not only by ARB treatment but ACEi as well (Fig 3.3A). That discrepancy may be due to mechanisms not captured by our model. It is possible that the improvement in bone mineral density predicted by our model for ACEi is not enough to have a notable impact on fracture risk. We also note that bone mineral density is not the only factor that impacts fracture risk; the architecture of the skeleton, alcohol usage, smoking, physical activity, and previous fractures are also major factors in fracture risk that are not represented in the model in this study [49].

### 4.1 Clinical perspectives

Although estrogen replacement therapy (or hormone replacement therapy) can effectively manage menopause symptoms and post-menopausal osteoporosis, its benefits must be weighed against the heightened risks of blood clots, stroke, and breast cancer, especially with long-term use [50, 51]. Furthermore, side effects such as weight gain, diarrhea, and mood changes can affect patient well-being and adherence [51]. The balance of risks and benefits associated with hormone replacement therapy varies considerably based on individual physiology, time since menopause onset, and the specific type of hormone replacement therapy [50]. Other current osteoporosis treatments, including antiresorptives and osteoanabolic therapies, have limitations on their duration of use, typically ranging from one to five years [52]. Therefore, the development of alternative long-term osteoporosis treatments are needed to broaden the range of effective options for patients. Our mathematical modeling analysis presented in this study supports clinical evidence suggesting that ARBs may contribute to maintaining bone health during post-menopausal estrogen decline by reducing AT1R- bound Ang II levels.

Indeed, there are a number of clinical applications that this model could be extended to study. The RAS is a key hormone system in blood pressure regulation. The RAS model used in this study is part of a larger model that involves whole-body blood pressure regulation [18, 19]. Using a similar approach to Ref. [53], the model presented in this study may be incorporated into the blood pressure model (Ref. [19]) through the RAS to investigate the interactions of hypertension and bone remodeling. Chronic kidney disease (CKD) is a disorder of the loss of kidney function that affects over 10% of the global population [54]. Many patients with CKD experience CKD-metabolic bone disorder, where their impaired renal function results in loss of bone mineral density. Previous mathematical modeling studies have used the underlying calcium homeostasis and bone remodeling model to study this disorder [15, 16, 55]. However, none have considered the RAS. Many CKD patients take RAS inhibitors to protect their kidney function, prevent hypertension, and have been shown to improve bone health [56]. The model presented in this study may be used to investigate how RAS inhibitors may impact bone density and electrolyte balance in patients with CKD.

### 4.2 Limitations and future directions

In this study, we developed a mathematical model that describes the interactions between the RAS, calciotropic hormones, calcium regulation, and bone remodeling. One major limitation of this model is the large number of unknown parameters, resulting in uncertainty in the model outcomes. In this study, we analyzed the impacts of variations in parameter values on model output by conducting a global sensitivity analysis using the Sobol method. However, we do recognize that there may be parameters and mechanisms that are not included or could be changed in this model.

A worthwhile future direction is to accurately model an aging population. In this study, we conducted simulations that represent the estrogen decline that is seen in post-menopausal women. However, aging populations, including post-menopausal women, have shown other physiological changes related to the systems in the model presented in this study. For example, aging populations have shown a blunted response to ACE inhibitors, have a lower GFR, decreased intestinal calcium absorption, lower vitamin D levels, and higher PTH levels [57–59]. These effects are not included in the model developed in this study at this time. Indeed, future work may consider the specific physiological impacts of aging for a more comprehensive description of post-menopausal physiology.

Given the importance of estrogen and using a female-specific RAS, the model presented in this study only represents a female population. However, sex differences have been found in the RAS [60] and renal handling of calcium [61, 62]. Men also can form osteoporosis as they age, which is not driven by the declining estrogen levels found in post-menopausal women. Rianon et al. [63] found that there are sex- and race-specific effects of ACEi on the prevention of age-related bone loss. Specifically, the authors reported that ACEi were effective at protecting against bone loss in African-American elderly men [63]. This is different than reports that did not consider sex and race differences and only reported significant changes in fracture risk in patients using ARB. The model presented in this study may be extended to consider how underlying sex and racial differences may explain altered treatment outcomes [64].

In this study, we do not include the hormone aldosterone, which is connected with the RAS as a part of the renin angiotensin aldosterone system. Researchers have shown that aldosterone also has a bidirectional relationship with the calciotropic hormones [6]. Clinical reports have shown that the drug spironolactone (an aldosterone inhibitor), has a significant impact on the bone health of hypertensive patients [65]. Indeed this hormone may be added to this model to study the impacts of hypertension, spironolactone, hyperaldosteronism, or other relevant conditions and medications on bone density and calcium regulation. Additionally, in this study, we assume that the RAS activity in the bone is the same as the systemic RAS, though this may not always be the case. Experimental evidence has shown that there is a bone-specific RAS [66]. Future work may involve modeling a bone-specific RAS to consider the impacts on bone remodeling similar to the renal-specific RAS models in Refs. [17, 67].

Estrogen levels vary during the different phases of the menstrual cycle in menstruating women [22]. Women who take oral contraceptives also have different estrogen levels [68]. The mathematical model presented here may be adapted to consider how changes in estrogen impacts the RAS, calcium homeostasis, and bone remodeling during the different phases of the menstrual cycle or on oral contraceptives [69]. During pregnancy, estrogen levels rise dramatically, while post-parturition, the estrogen levels drop below pre-pregnancy levels to support lactation [70]. Each of these changes results in major impacts on physiology, including the RAS, calcium homeostasis, and bone remodeling [70, 71]. Previously, we developed mathematical models that represent calcium homeostasis in rats that consider sex differences and maternal adaptations of pregnancy and lactation in Ref. [72], but these models do not include a detailed bone compartment or the impact of the RAS. The model presented in this study may be similarly adapted to consider the major changes that women undergo during pregnancy and lactation.

## Grants

This work is supported by the Canada 150 Research Chair program and by the National Science and Engineering Research Council of Canada via a Discovery award (RGPIN-2019-03916) to A.T.L. and a Canada Graduate Scholarship (CGS-D) to M.M.S. The funders had no role in study design, data collection and analysis, decision to publish, or preparation of the manuscript.

## Author Contributions

A.T.L. and M.M.S. conceived and designed research; M.M.S. wrote model code; M.M.S. conducted simulations; M.M.S. and A.T.L. analyzed data; M.M.S. and A.T.L. interpreted results of simulations; M.M.S. prepared figures; M.M.S. and A.T.L. drafted manuscript; M.M.S. and A.T.L. edited and revised manuscript; M.M.S. and A.T.L. approved final version of manuscript.

## Supplemental Materials

### A Impact of declining estrogen levels on the RAS

How do the various impacts of estrogen on the RAS change RAS levels during post-menopausal estrogen decline? To investigate this question, we conducted a simulations of the estrogen-RAS model presented in this study. The results are shown in Fig. A.1. We assume that estrogen decline starts at age 50 years (Fig. A.1A; Eq. 2.1). The decline in estrogen levels, decreases AGT production so that by age 80 years [AGT] is 60% below pre-menopausal levels (Fig. A.1B). Plasma renin secretion is inhibited by estrogen levels, which are rapidly decreasing. Renin secretion is inhibited by estrogen, so the lowered estrogen levels result in driving increased renin secretion so that ultimately plasma renin levels rise (Fig. A.1C).

**Figure A.1.**
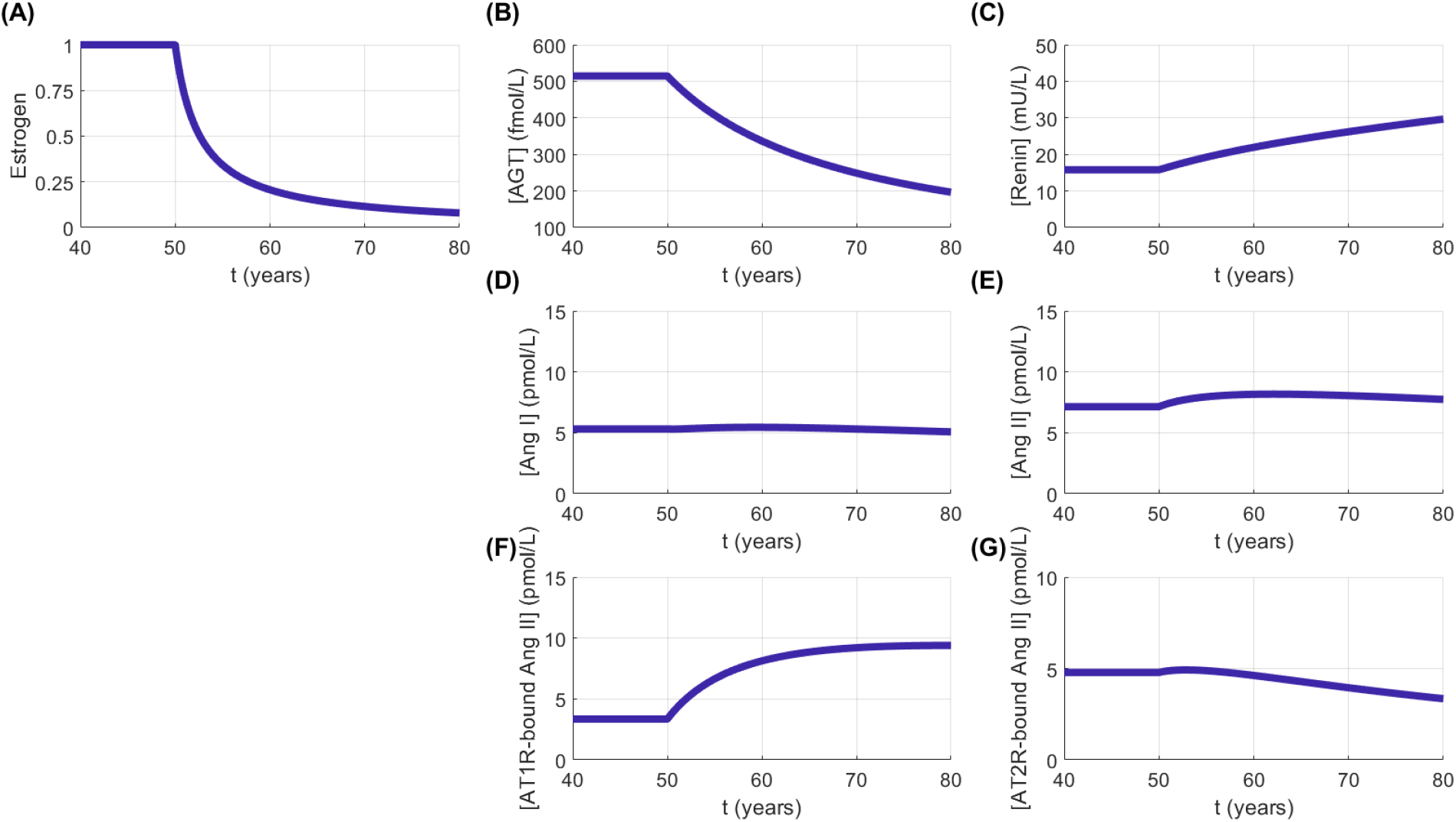
Simulation results of the impact of decreasing (A) estrogen levels on the plasma concentration of (B) AGT, (C) renin, (D) Ang I, (E) Ang II, (F) AT1R-bound Ang II, and (G) AT2R-bound Ang II. AGT: angiotensinogen, Ang: angiotensin, AT1R: angiotensin type 1 receptor, AT2R: angiotensin type 2 receptor

While renin concentration increases, the decreased AGT levels result in decreased PRA, which drives Ang I production. This results in stable Ang I levels (Fig. A.1D). ACE activity is increased so we see a slight increase in Ang II levels (Fig. A.1E). Ang II then binds to AT1R and AT2R. Estrogen inhibits AT1R binding and activates AT2R binding (Fig. 2.1). When estrogen levels decrease, AT1R receptors are inhibited less, thus leading to increased AT1R-bound Ang II levels (Fig A.1F). For AT2R receptors, initially, AT2R-bound Ang II levels slightly increase from baseline due to the increased Ang II levels, but later AT2R-bound Ang II levels decrease from baseline due to the loss of activation by estrogen levels (Fig. A.1G).

We conducted a Sobol sensitivity analysis on the estrogen-RAS model to investigate which parameters impact the estrogen deficiency model simulation the most for AGT, Ang II, AT1R-bound Ang II, and AT2R-bound Ang II. We varied the parameters that represent the rate of enzyme impacts on the RAS (i.e., parameters *c*_*i*_) as well as the parameters that represent estrogen impact on the RAS (i.e., parameters *e*_*i*_ listed in Table 2.1). Each parameter was varied by a 2-fold increase and decrease from the baseline value. We then conducted a Sobol analysis at the time point of every 10 years for the estrogen deficiency simulation.

The results for the total order Sobol indices (*S*_total_) over the estrogen decline RAS simulation are shown in Fig. A.2. Before the onset of estrogen decline due to menopause (i.e., *t <* 50), the parameters *e*_*i*_ (*i*= renin, AGT, ACE, etc), which represent the impact of estrogen on the RAS (Eqs. 2.2–2.3), do not have any impact on the model variation as expected because *E* = 1.

**Figure A.2.**
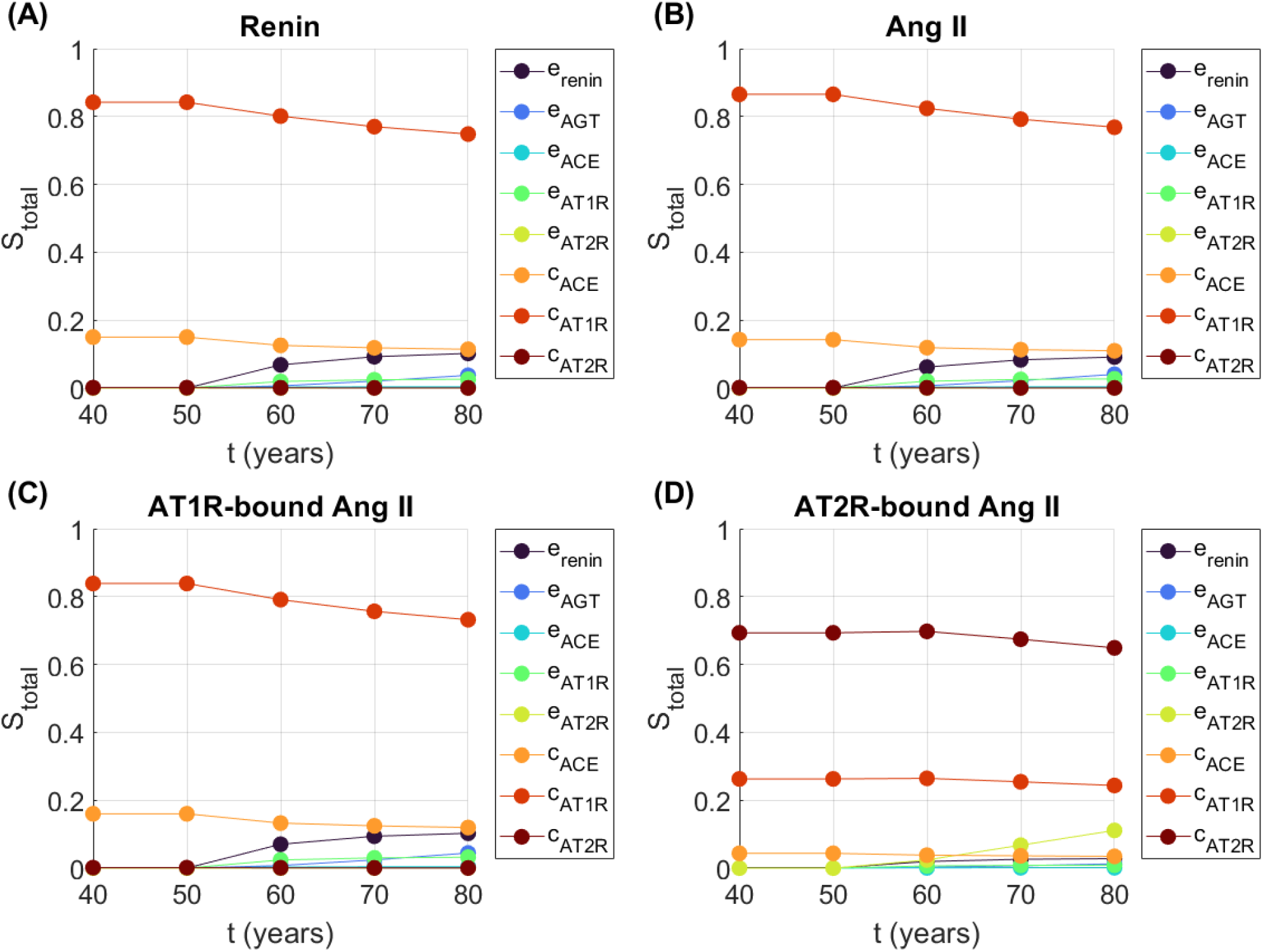
Sobol total indices (S_total_) for estrogen impact on (A) AGT, (B) Ang II, (C) AT1R-bound Ang II, and (D) AT2R-bound Ang II at different time points in the simulation. Parameters not shown did not have a S_total_ over 0.01 for any variable. AGT: angiotensinogen, Ang: angiotensin, AT1R, angiotensin type 1 receptor, AT2R: angiotensin type 2 receptor

For renin, *e*_renin_ represents the impact of estrogen on renin secretion and leads to about 10% of the model output variation by *t* = 70 years (Fig. A.2A). The parameters with the largest output on model output of Ang II are *c*_ACE_ and *c*_AT1R_ which correspond to ACE activity and AT1R binding, respectively (Fig. A.2B). After the decline in estrogen, the impact of estrogen on renin secretion, *e*_renin_ contributes to the variation in the model output of Ang II as well (Fig. A.2B). AT1R-bound Ang II is also impacted by *c*_ACE_ and *c*_AT1R_ the most due to the dependence on Ang II levels (Fig. A.2C). After the onset of estrogen decline, the impact of estrogen on renin and AT1R binding contribute to variation in the model output. By the end of the simulation time, about 10.2% and 3.2% of model output variation of AT1R-bound Ang II can be attributed to *e*_renin_ and *e*_AT1R_, respectively (Fig. A.2C). For AT2R-bound Ang II, about 11% of the total variation can be attributed to the impact of estrogen on AT2R binding, *e*_AT2R_, by the end of the simulation time (Fig. A.2D).

### B Additional Figures

**Figure B.1.**
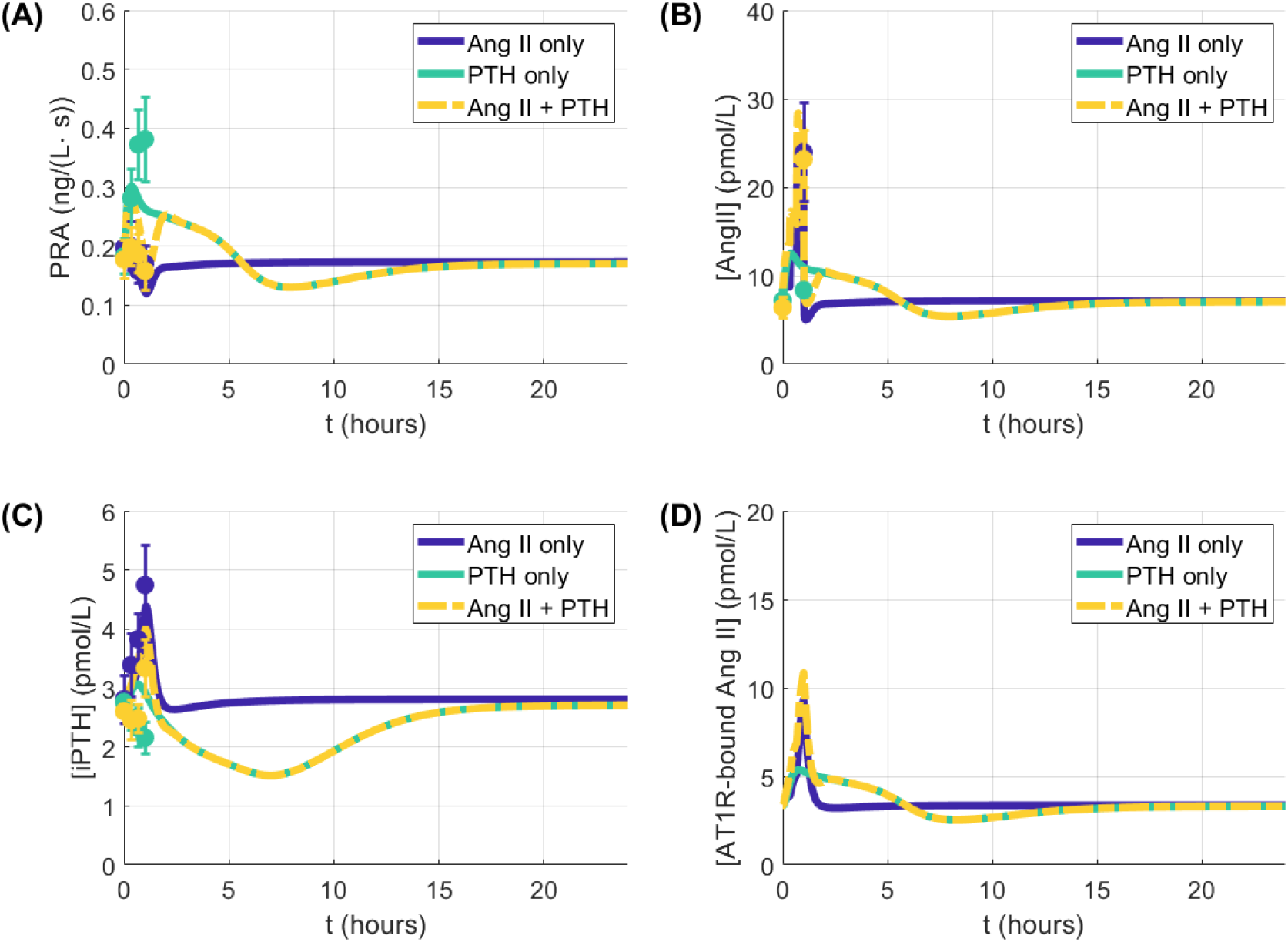
Replication of Fig. 3.1 for a longer simulation time. Simulation results for replication of the experiments in Grant et al. [37]. Results for (A) plasma renin activity (PRA), (B) angiotensin II (Ang II) plasma concentration, (C), parathyroid hormone (PTH) concentration, and (D) angiotensin receptor type II bound Ang II are shown for during and after 1 hour of infusions of Ang II only, PTH only, and Ang II and PTH infusions together. Lines indicate simulations, while the dots and error bars in panels A–C are from the relevant experimental data presented in Grant et al. [37]. AT1R: angiotensin receptor type 1, iPTH: intact parathyroid hormone

**Figure B.2.**
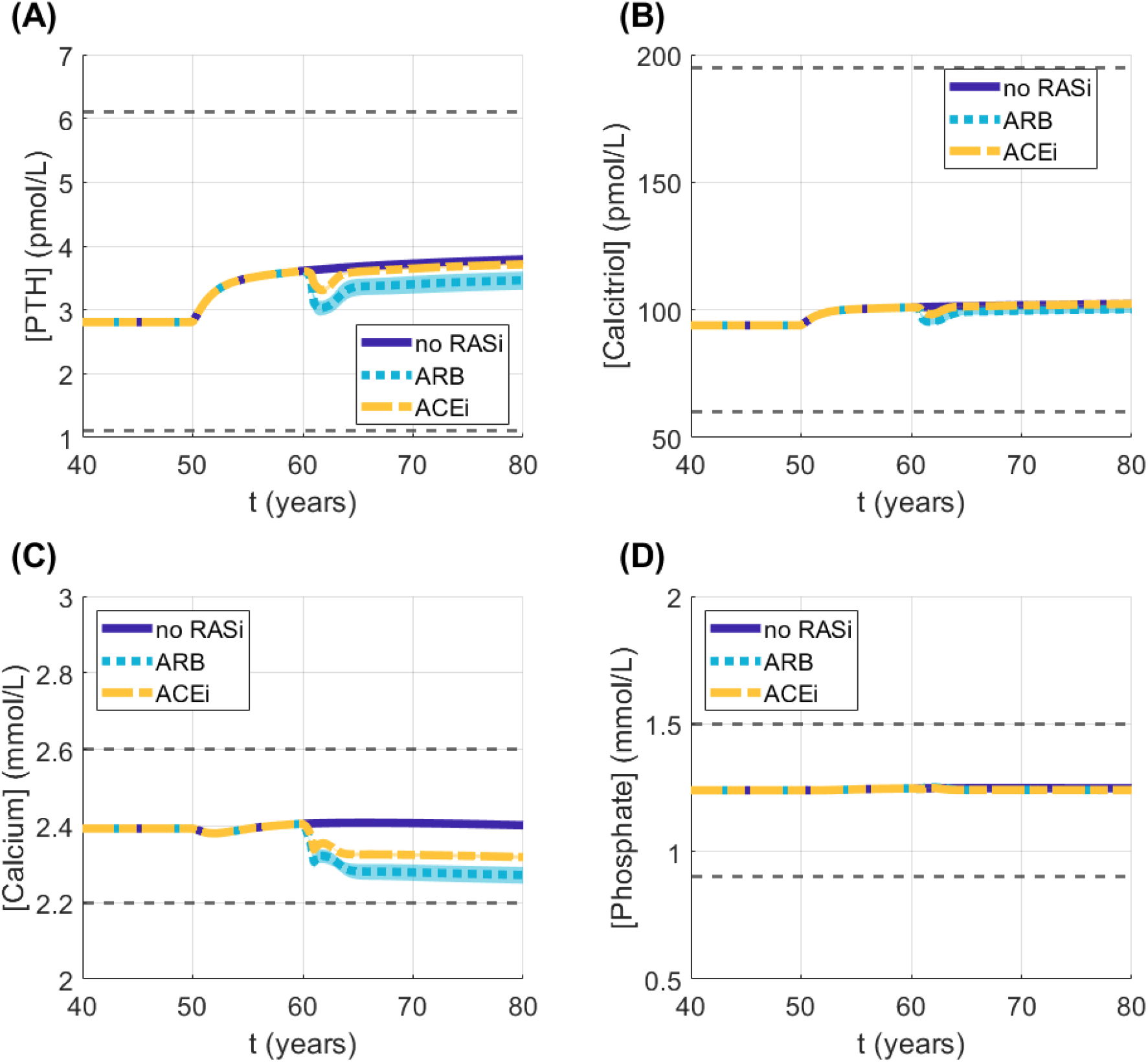
Impact of estrogen deficiency and RAS inhibitors on extracellular levels of calciotropic hormones (A) parathyroid hormone and (B) calcitriol as well as electrolytes (C) calcium and (D) phosphate. For the angiotensin receptor blocker simulations (labeled “ARB” in the legend), σ_ARB_ was varied between 92–98.3% based on the ranges reported in Ref. [18]. For the angiotensin-converting enzyme inhibitor simulations (labeled “ACEi” in the legend), σ_ACEi_ was varied between 94–97.2% as reported in Ref. [18]. Lines represent the mean values over the varied σ_i_ values for the respective treatment with shaded area representing the range. PTH: parathyroid hormone, RASi: RAS inhibitor, ARB: angiotensin receptor blocker, ACEi: angiotensin-converting enzyme inhibitor

### C Key parameter descriptions

**Table C.1:** Descriptions of parameters shown in Figs. 3.5, 3.6, and 3.7. The reference is the original source of the parameter value. Ang II: angiotensin II, AOH: 1α-hydroxylase, AT1R: angiotensin type 1 receptor, CaSR: calcium sensing receptor, OC: osteoclast, PTH: parathyroid hormone

| Name | Description | Reference |
| --- | --- | --- |
| $E_{\max,L-PTH}$ | half-maximal response of effect of PTH on RANKL production | [13] |
| $k_3$ | binding rate of RANK and RANKL | [13] |
| $k_4$ | unbinding rate of RANK-RANKL | [13] |
| $k_{in,L}$ | baseline RANKL production | [13] |
| $k_{out,L}$ | degradation rate of RANKL | [13] |
| $k_{in,RNK}$ | baseline RANK production | [13] |
| $k_{out,RNK}$ | degradation rate of RANK | [13] |
| $\rho_{AT1R,L}$ | minimum impact of AT1R on RANKL | Eq. 2.22 |
| $\delta_{AT1R,L}$ | half-maximal response of AT1R impact on RANKL | Eq. 2.22 |
| $\alpha_{CaSR,renin}$ | maximum impact of CaSR on renin | Eq. 2.18 |
| $\delta_{CaSR,renin}$ | half-maximal response of CaSR impact on renin | Eq. 2.18 |
| $\Phi_{Plas2BoneCa0}$ | baseline calcium exchange from extracellular fluid to bone | [13] |
| $\Phi_{Bone2PlasCa0}$ | baseline calcium exchange from bone to extracellular fluid | [13] |
| $c_{AT1R}$ | binding rate of Ang II to AT1R | [18, 19], Eq. 2.10 |
| $N_{rs}$ | baseline renin secretion | [18, 19], Eq. 2.4 |
| $X_{PRCPRA}$ | equilibrium ratio of PRA to PRA | [18] |
| $\delta_{PTH,renin}$ | half-maximal response of PTH impact on renin | Eq. 2.19 |
| $K_{Ca,PTH}$ | half-maximal response of PTH secretion to CaSR | [13] |
| $k_{base,AOH}$ | baseline AOH production | [13] |
| $k_{deg,AOH}$ | degradation rate of AOH | [13] |
| $V_{\max,PTH-AOH}$ | maximum effect of PTH on AOH production | [13] |
| $\delta_{AOH,Phos}$ | half-maximal response of impact of phosphate on AOH | [13] |
| $V_{\max,Phos}$ | maximum impact of phosphate on AOH production | [13] |
| $V_{\max,PTH-reabs}$ | maximum impact of PTH on renal reabsorption | [13] |

### D Model equations

A list of the ordinary differential equations in the model is given below.

#### D.1 Renin-angiotensin system

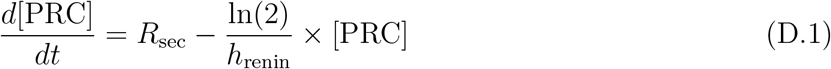

where

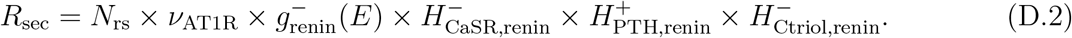

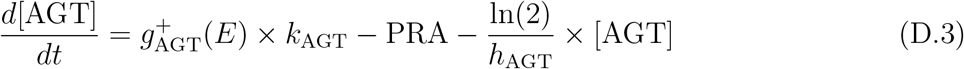

where

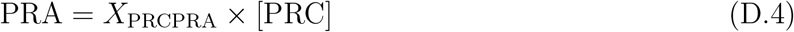

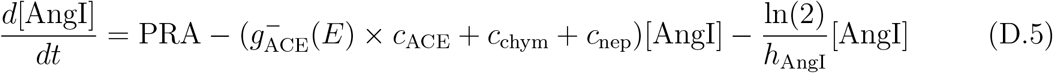

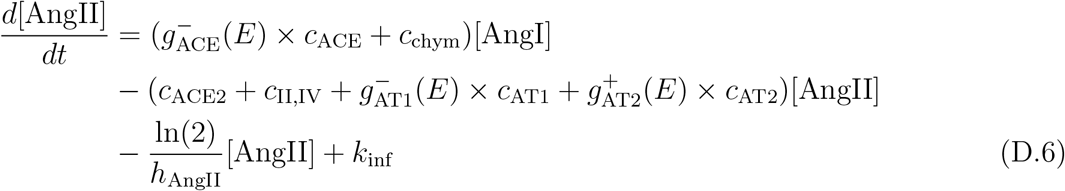

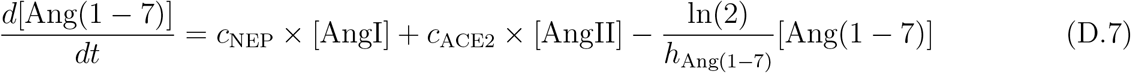

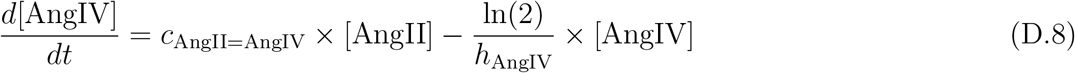

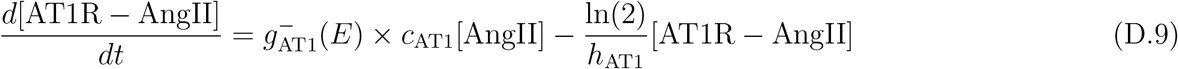

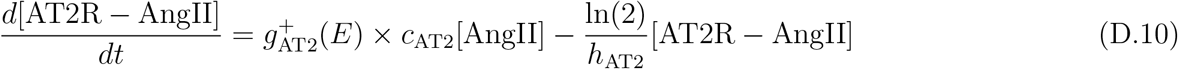

#### D.2 Calcium regulation

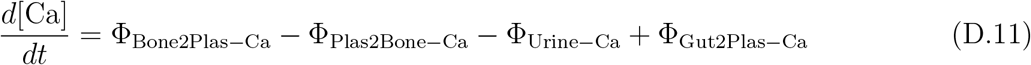

with

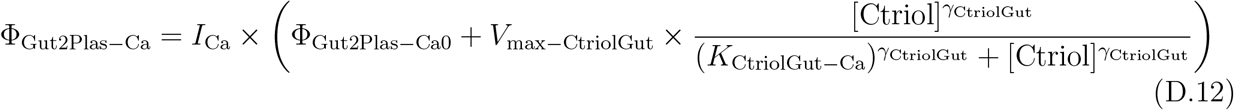

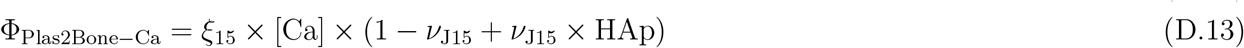

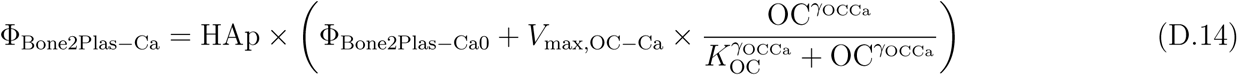

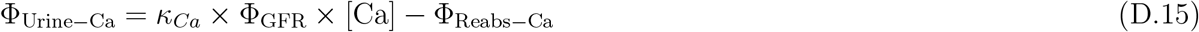

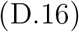

where

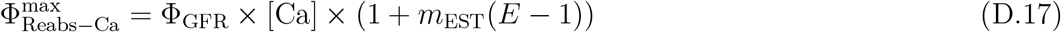

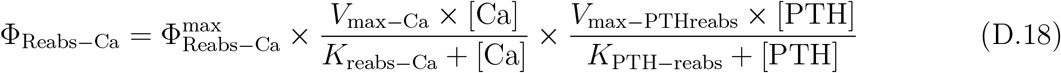

#### D.3 Parathyroid hormone

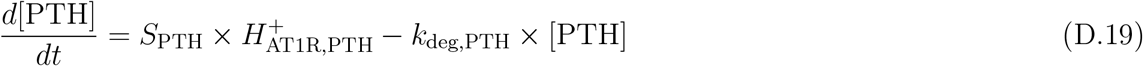

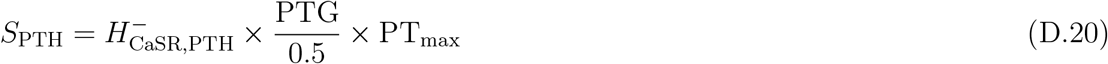

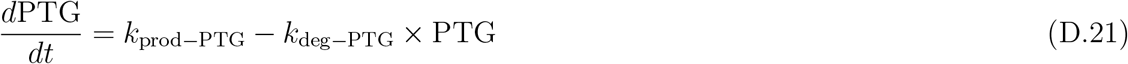

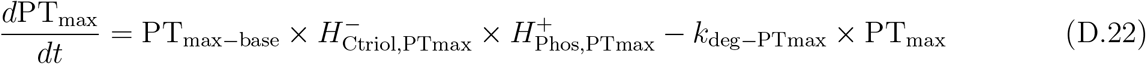

#### D.4 Calcitriol

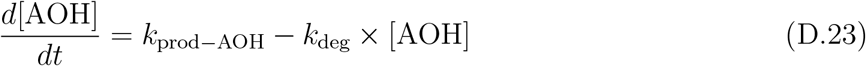

with

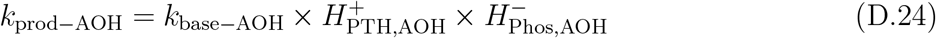

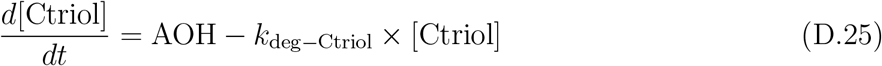

#### D.5 Phosphate regulation

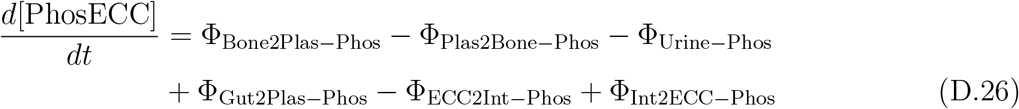

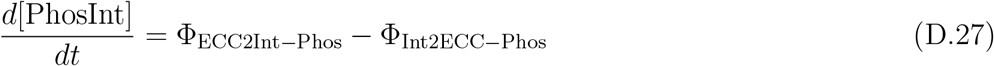

with

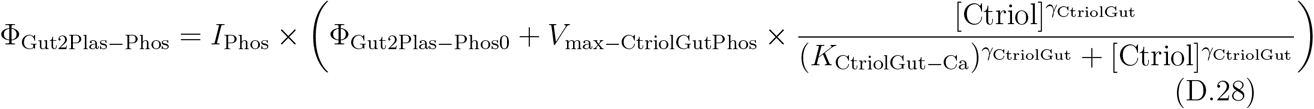

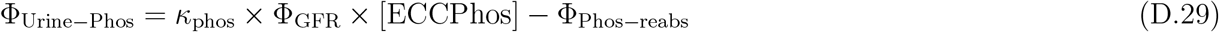

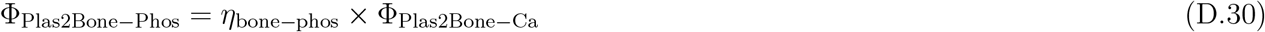

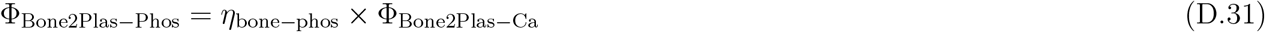

where

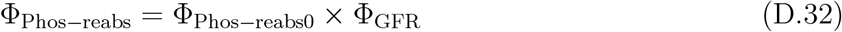

#### D.6 Bone remodeling

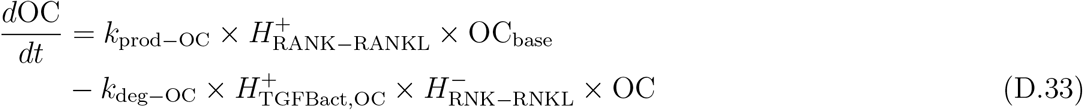

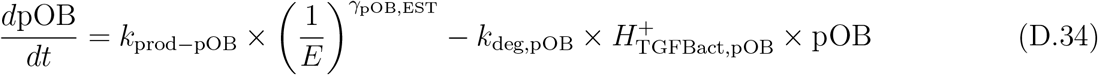

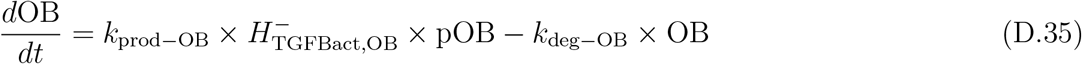

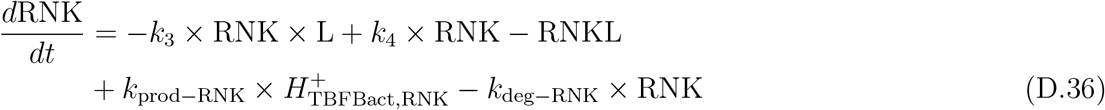

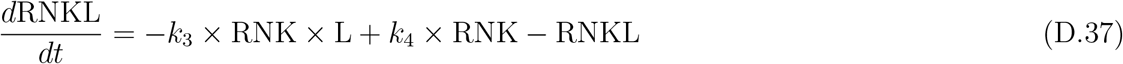

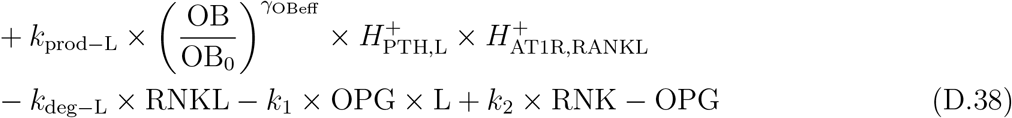

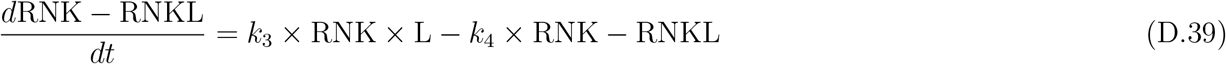

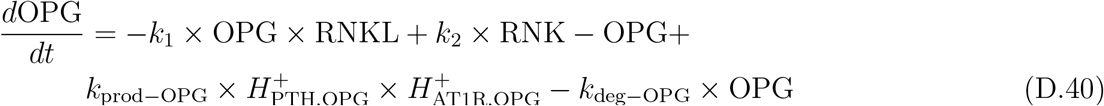

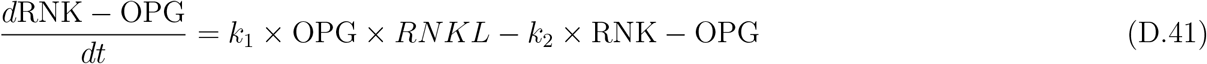

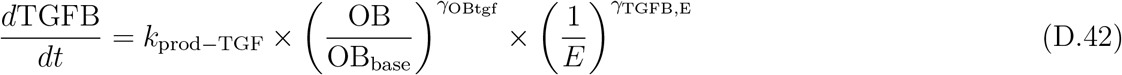

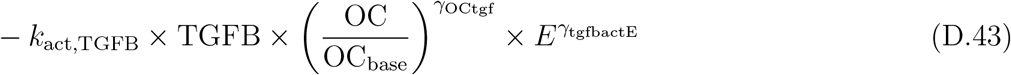

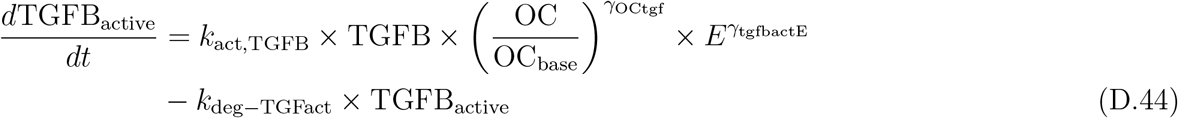

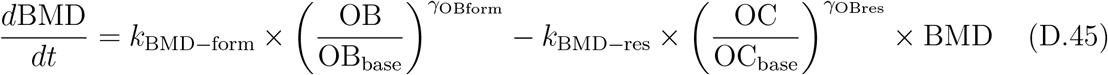

##### D.7 List of variable names

AGT: angiotensinogen
Ang I: angiotensin I
Ang II: angiotensin II
Ang (1-7): angiotensin (1-7)
Ang IV: angiotensin IV
AOH: 1*α*-hydroxylase
AT1R-AngII: angiotensin type 1 receptor bound Ang II
AT2R-AngII: angiotensin type 2 receptor bound Ang II
BMD: bone mineral density
Ca: plasma calcium
Ctriol: calcitriol in plasma
*E*: normalized estrogen level
OB: osteoblasts
OC: osteoclasts
OPG: osteoprotegerin
PhosECC: extracellular phosphate
PhosInt: intracellular phosphate
pOB: pre-osteoblasts
PRA: plasma renin activity
PRC: plasma renin concentration
PT_max_: maximal parathyroid gland secretion
PTG: parathyroid hormone in the parathyroid glands
PTH: parathyroid hormone
RNK: RANK
RNKL: RANKL
TGFB: latent TGF*β*
TGFB_active_: active TGF*β*

